# Across-population genomic prediction in grapevine opens up promising prospects for breeding

**DOI:** 10.1101/2021.07.29.454290

**Authors:** Charlotte Brault, Vincent Segura, Patrice This, Loïc Le Cunff, Timothée Flutre, Pierre François, Thierry Pons, Jean-Pierre Péros, Agnès Doligez

**Author notes:** Corresponding author: Agnès Doligez, UMR AGAP Institut, Univ Montpellier, CIRAD, INRAE, Institut Agro, F-34398 Montpellier, France.

## Abstract

Crop breeding involves two selection steps: choosing progenitors and selecting offspring within progenies. Genomic prediction, based on genome-wide marker estimation of genetic values, could facilitate these steps. However, its potential usefulness in grapevine (*Vitis vinifera* L.) has only been evaluated in non-breeding contexts mainly through cross-validation within a single population. We tested across-population genomic prediction in a more realistic breeding configuration, from a diversity panel to ten bi-parental crosses connected within a half-diallel mating design. Prediction quality was evaluated over 15 traits of interest (related to yield, berry composition, phenology and vigour), for both the average genetic value of each cross (cross mean) and the genetic values of individuals within each cross (individual values). Genomic prediction in these conditions was found useful: for cross mean, average per-trait predictive ability was 0.6, while per-cross predictive ability was halved on average, but reached a maximum of 0.7. Mean predictive ability for individual values within crosses was 0.26, about half the within-half-diallel value taken as a reference. For some traits and/or crosses, these across-population predictive ability values are promising for implementing genomic selection in grapevine breeding. This study also provided key insights on variables affecting predictive ability. Per-cross predictive ability was well predicted by genetic distance between parents and when this predictive ability was below 0.6, it was improved by training set optimization. For individual values, predictive ability mostly depended on trait-related variables (magnitude of the cross effect and heritability). These results will greatly help designing grapevine breeding programs assisted by genomic prediction.

## Introduction

Breeding for perennial species is mostly based on phenotypic selection and is hindered by cumbersome field trials and the long generation time. Genomic prediction (GP), based on genome-wide prediction of genetic values ^1^, has been widely adopted in modern plant and animal breeding programs, for its superiority in terms of cost and time saved compared to traditional phenotypic selection, but also because it allows handling traits with complex genetic determinism. GP requires a model training step within a reference population, prior to model application to a target population of selection candidates ^2^. In perennial crops, a universal population encompassing most of the species’ genetic diversity could be particularly interesting as a training population to reduce phenotyping effort, since breeding cycle and juvenile phase are long.

Breeding schemes typically involve first the choice of parents (individuals to be crossed) and then the selection of offspring within crosses. GP is adapted both for predicting cross mean and for ranking genotypes within a cross (Mendelian sampling). These steps correspond to the components of the predictive ability (PA) of GP. It is indeed defined as the sum of cross mean and Mendelian sampling terms, as detailed in Werner et al. ^3^.

Under an additive framework, cross mean is expected to be the sum of the breeding values of parents, but some deviation may result from dominance or epistasis ^4^. So far, a few studies only have investigated cross mean PA ^5, 6, 7, 8^, although none of them clearly investigated its influencing parameters.

In contrast, the prediction of genetic values within a cross (Mendelian sampling), has been widely studied, both with simulated and real data. Various parameters affecting PA have been pointed out, including the statistical method used ^9^, the composition and size of training and validation populations ^10, 11^, the trait genetic architecture and heritability ^12, 13^ and marker density ^14^. Genetic relationship between the training and validation sets is known to strongly affect PA ^15^, with low or even sometimes negative accuracies for across-breed GP in animals ^16^. This can be explained by the loss of linkage phase between the marker and QTL or by differences in linkage disequilibrium among populations ^17^. Another explanation is the presence of specific allelic effects and allele frequencies, due to the genetic background ^18^. All these effects are linked to genetic relationship. Some studies specifically derived deterministic equations to predict PA for across-population GP, based on genetic relationship and heritability (e.g., 19, 20, 21).

In grapevine (*Vitis vinifera* subsp. *vinifera*), very few authors have assessed the potential interest of GP. Viana et al. ^22^ investigated GP within a bi-parental population from a cross between an interspecific hybrid and a seedless table grape. Later, Migicovsky et al. ^23^ used a panel of 580 *V.vinifera* accessions to perform both GP and genome-wide association study (GWAS) for 33 phenotypes. More recently, Brault et al. ^24^ investigated GP within a bi-parental population from a cross between Syrah and Grenache. In a related study, Fodor et al.^25^ had simulated a structured and highly diverse grapevine panel and bi-parental populations with parents originating from the panel. They applied GP and found little difference between PA values estimated within the panel or across populations. Finally, Flutre et al. ^26^ studied 127 traits with GWAS and GP within a diversity panel; they also applied across-population GP, but with 23 test offspring and for one trait only. Before genomic selection can be deployed in grapevine, evaluating PA across populations is thus crucially needed. In particular, PA should be evaluated with a diversity panel and a bi-parental progeny as training and validation sets, respectively, a configuration much more likely to occur in actual breeding schemes than GP within the same population. As in grape, studies investigating across-population GP are also lacking in most clonally propagated crops.

The aim of this study was to assess across-population genomic PA and to provide a more thorough understanding of parameters affecting PA in a situation close to the one typically encountered in a breeding context, i.e. across populations, for a clonally propagated crop such as grapevine. Our study was based on phenotypic data for 15 traits, related to yield, berry composition, phenology and vigour, measured both in a diversity panel ^27^, and in a half-diallel with 10 bi-parental crosses. We assessed PA under three scenarios, first for cross mean, and then for Mendelian sampling term; the results provided keys to understand PA determinants in both cases. Finally, we implemented training population optimization to investigate under which conditions PA can be improved.

## Results

### Extent of genetic diversity within the half-diallel population

We first evaluated the genetic variability of half-diallel crosses with respect to the diversity panel, through their projection on the first plane of a PCA based on genotypic data at 32,894 SNPs within the diversity panel. The half-diallel crosses were genetically close to the wine west (WW) subpopulation from the diversity panel (Figure 1a), which was expected, given that all half-diallel parents except Grenache are wine varieties from western Europe (Figure 1a, Figure S1). The half-diallel diversity covered the whole range of WW diversity, and progenies, all located exactly between their respective parents, were well separated from each other along the first two PCA axes (Figure 1a).

**Figure 1:**
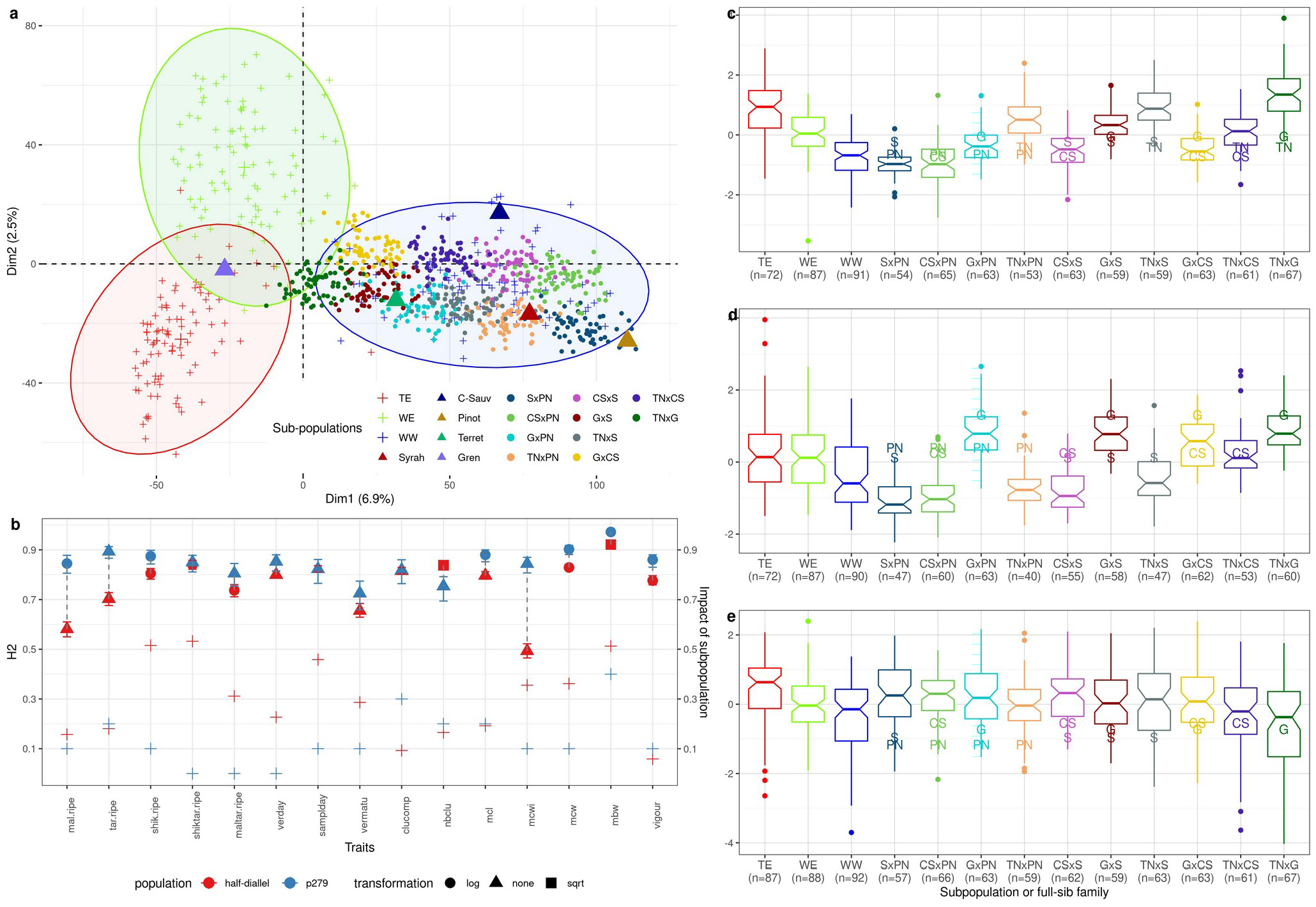
Description of the half-diallel, relative to the diversity panel. a: PCA of the diversity panel based on 32,894 SNPs with the 3 subpopulations distinguished by different colors, on which half-diallel progenies (dots) and parents (triangles) were projected. b: Broad-sense heritability estimates in the whole half-diallel (red) and in the diversity panel (blue) for the 15 traits studied (left axis), with shape corresponding to the transformation applied to raw data; the relative variance due to the cross effect and the R^2^ of the subpopulation effect, for the half-diallel (red) and the diversity panel (blue), respectively, are also reported with ‘+’ (right axis). c, d, e: genotypic value BLUP distribution in each subpopulation or progeny, for mean berry weight, mean cluster width and vigour, respectively; BLUPs for parents are indicated by their initial letters (Table S4). Number of genotypes per subpopulation/progeny is indicated below the subpopulation/progeny name. These traits were chosen to represent various levels of H^2^ and relative importance of cross effect. BLUP distributions for all traits are presented in Figure S3.

We then investigated broad-sense heritability values (H^2^) for 15 traits related to yield, berry composition, phenology and vigour. Overall H2 values were medium to high, ranging from 0.49 for **mcwi** in the half-diallel to 0.92 for **mbw** in the panel (Figure 1b; Table S1). Correlation between half-diallel and diversity panel heritability values was 0.31. Per-cross H^2^ values for each trait varied among half-diallel crosses (Figure S2), which might result from the fairly small number of offspring per cross (from 64 to 70). Nevertheless, we observed a 0.68 correlation between overall and per-cross H^2^. Mean cluster width displayed extreme variation in H^2^ per cross (from 0.02 to 0.67). This might be due to the difficulty to phenotype this specific trait because of the presence of lateral wings in some individuals.

Within the half-diallel and for all traits, the *cross* effect was retained in the mixed model for genetic value estimation, but its magnitude with respect to the total genetic variance varied depending on the trait, ranging from less than 10% to ca. 50% (Figure 1b; Table S1). Depending on the trait or cross, the distribution of genotypic BLUPs varied widely (Figure 1c-e; Figure S3), some traits such as **vigour** being quite comparable among crosses, while others such as **mbw** or **mcwi** varied greatly. We also observed transgressive segregation within the half-diallel progenies (Figure 1c-e; Figure S3) for most traits and subpopulations. The 15 traits studied represented a large phenotypic diversity, structured among crosses (Figure S4).

### Prediction of cross mean and Mendelian sampling within- and across-populations

We first implemented cross mean prediction, as if aiming to select parents for future crosses, selecting the method best adapted to genetic architecture between RR and LASSO (see Material and Methods). Predictive ability (PA) was assessed as Pearson’s correlation between the observed mean genotypic value per half-diallel cross and the one predicted based on parental average genotypes (Table S2). Three scenarios were tested (Material and Methods, Figure 2): allelic effects estimated within the whole half-diallel (scenario 1a), in families with one parent in common (scenario 1b), or within the whole diversity panel (scenario 2).

**Figure 2:**
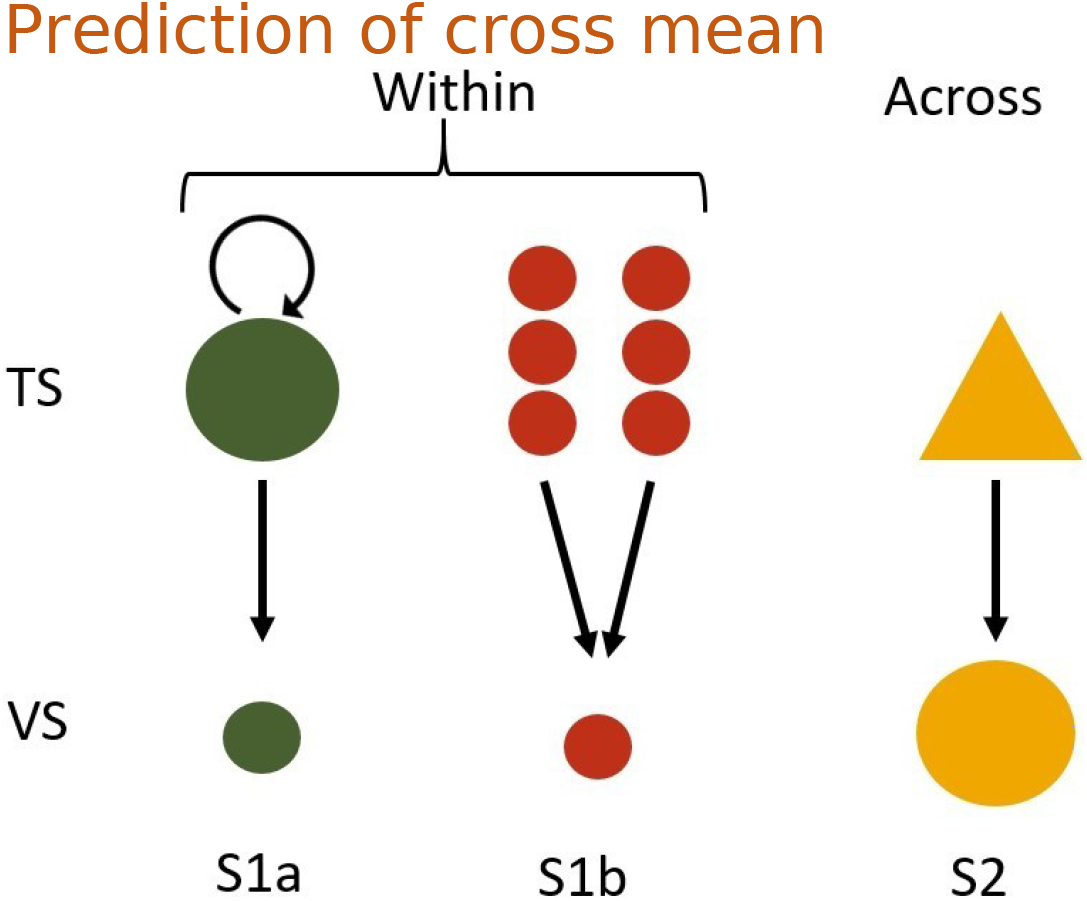
Schematic description of the three scenarios tested. TS: training set, VS: validation set. In scenario 1a, GP was applied within the half-diallel population with 10-fold cross-validation repeated 10 times. In scenario 1b, half-sib families from each parent were used separately as TS. In scenario 2, TS was the diversity panel. See details in Table S5.

In scenario 2, per-trait and per-cross predictive ability was lower and more variable than in scenarios 1a and 1b (Figure 3). Average per-cross PA was 0.56, 0.62 and 0.29 in scenarios 1a, 1b and 2, respectively (Figure 3a). Average per-trait PA was close to 1 for most traits in scenarios 1a and 1b (Figure 3b), and still high (around 0.75) in scenario 2, when excluding **nbclu** and **vigour** (Table S3). Overall PA (over the 150 cross x trait combinations) was 0.32. There was upward or downward bias for some traits, scenarios or methods, and in scenario 1a, LASSO resulted in larger bias (Figure S5).

**Figure 3:**
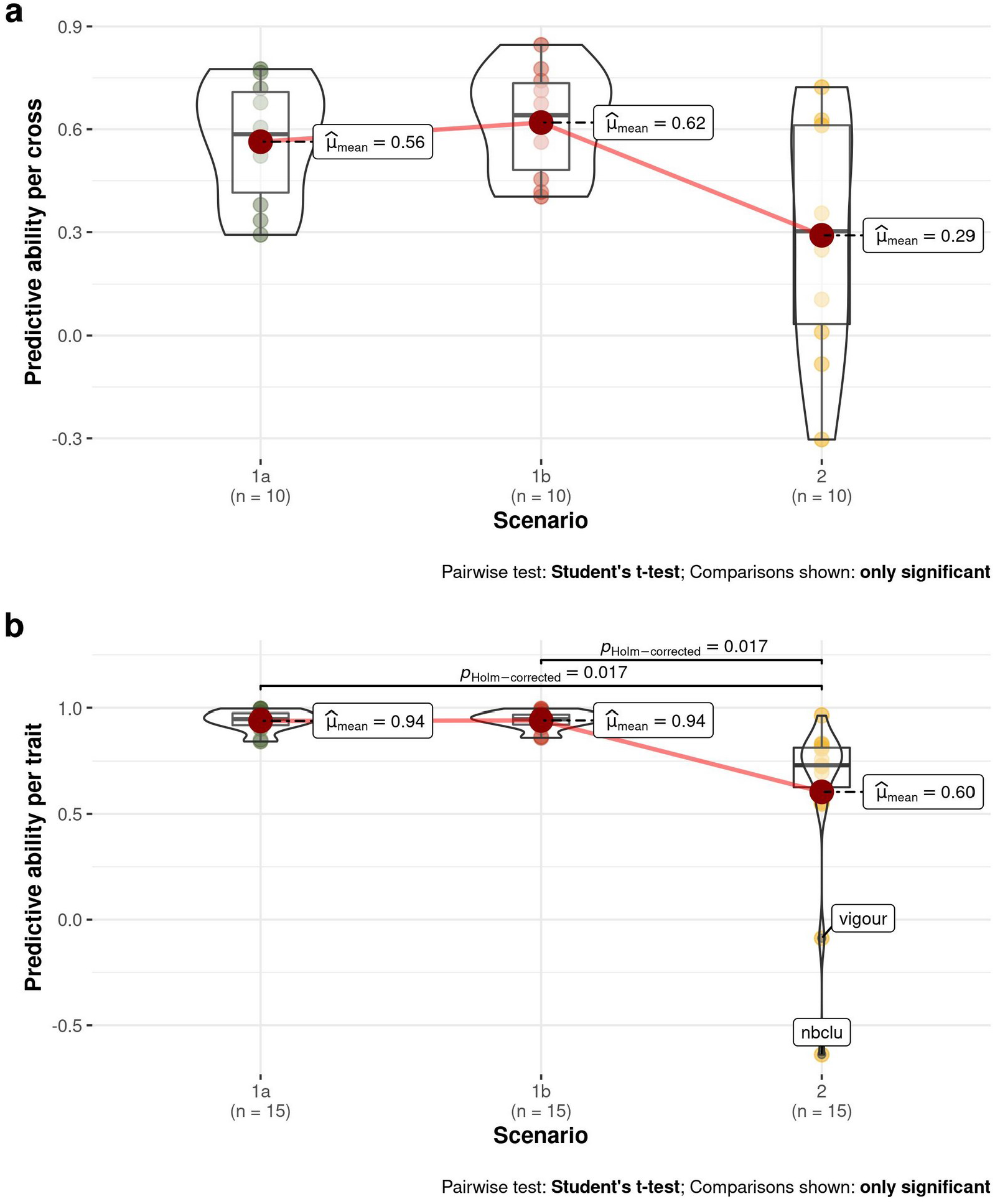
Boxplots of PA values for the three scenarios (1a: within whole half-diallel prediction; 1b: half-sib prediction within half-diallel; 2: across-population prediction with diversity panel as training set and each half-diallel cross as validation set). Each PA value was the best one obtained between RR and LASSO methods. Average PA is indicated next to each boxplot. a: per-cross PA, b: per-trait PA.

### Prediction of Mendelian sampling

We then measured PA for individual offspring within each half-diallel cross, thus considering separately the Mendelian sampling component. For each cross and trait, we compared the observed and predicted genotypic values in the three scenarios (Figure 2; Figure S6)

In scenario 1a (Figure 4a), average PA per trait ranged from 0.18 for **vermatu** to 0.58 for **mbw**, with a 0.47 overall average (Figure S7a). The extent of PA variation among crosses depended on the trait and could be very large, as for **vermatu** (from −0.074 to 0.443). Unlike for traits, no cross had constantly high or low PA (Figure S7b). RR method yielded the highest PA values in most cases (91% of the 150 trait x cross combinations).

**Figure 4:**
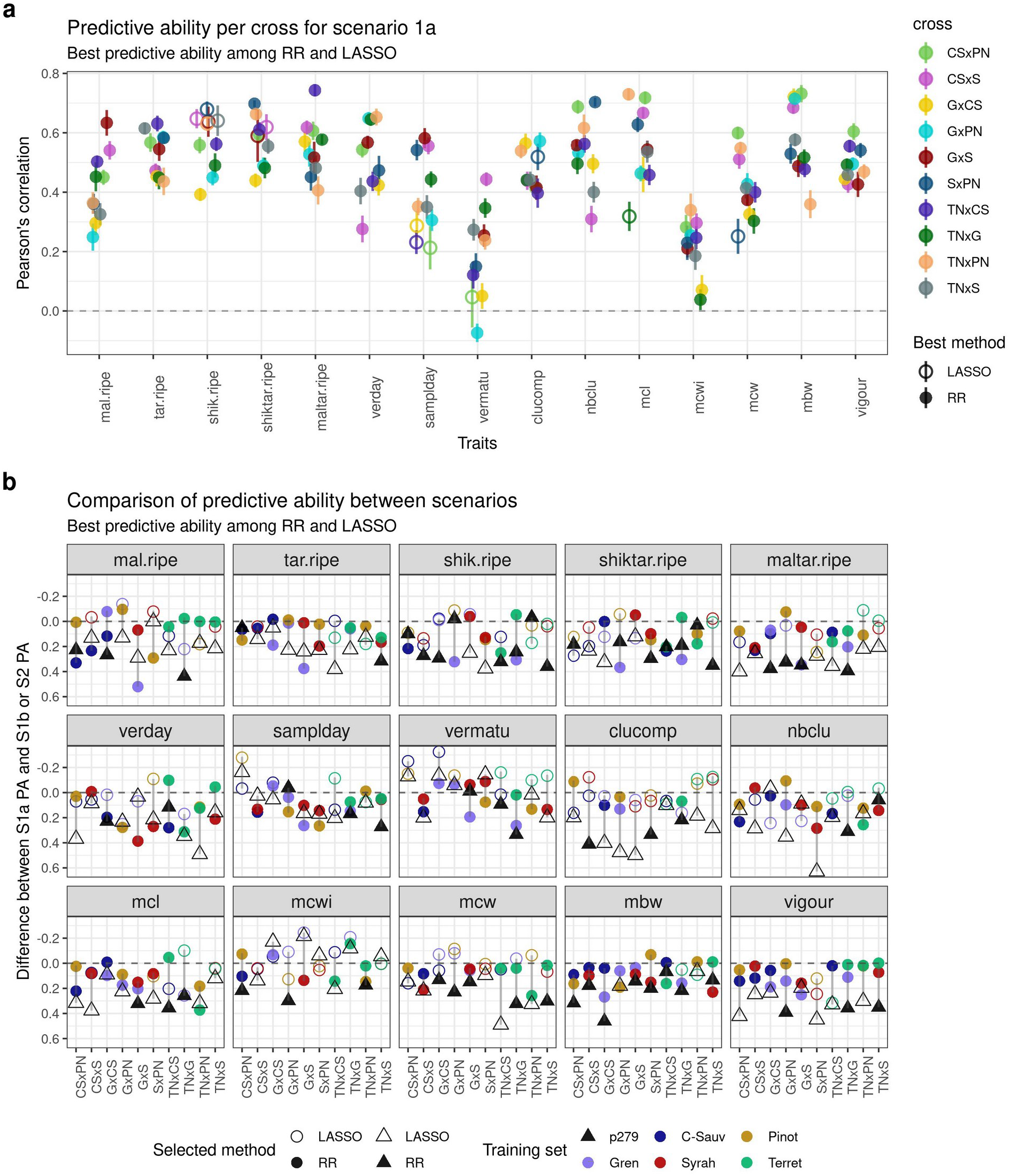
a: Mendelian sampling PA per trait and cross for scenario 1a with the best method between RR and LASSO. Vertical bars represent the standard error around the mean (95 % of the confidence interval), based on the outer cross-validation replicates. PA corresponds to the Pearson’s correlation between the BLUPs of the genotypic value and the predicted genotypic values. b: Difference between PA of scenario 1a and of the other scenarios. S2 is displayed with a triangle, and S1b by circles, colored according to the parental training set and filled if the best method was RR and empty otherwise.

In scenario 1b (Figure 4b), there were two PA values per cross, one for each parental training set (TS). The difference between these two values varied widely, depending on the cross and trait (up to about 0.5 for **mal.ripe** in GxS), with an overall average of 0.39. Most often, PA was lower in scenario 1b than in scenario 1a, likely because no full-sibs were included in the training set. However, there were several cases with PA values similar or higher in scenario 1b for one parental TS compared to scenario 1a. RR method produced the best PA in 61% of the 300 combinations (2 parents × 15 traits × 10 crosses).

In scenario 2 (Figure 4b), overall average PA (0.26) was nearly halved compared to scenario 1a, with trait dependent differences in PA between both scenarios. Some traits such as **vigour**, **clucomp** and **maltar.ripe** displayed a particularly marked decrease. On the opposite, **mcwi** and **vermatu** reached equivalent PA values in both scenarios. RR provided the best PA in 61% of the 150 combinations.

### Exploring factors affecting predictive ability, and training set optimization

We sought those variables affecting the PA values observed above, both for prediction of cross mean and Mendelian sampling. We then implemented training set (TS) optimization in an attempt to increase PA.

#### Variables affecting the prediction of cross mean

In scenario 2, per-cross PA was highly negatively correlated (−0.9) with the cross parents’ pairwise distance on the first axis of the diversity panel PCA (Figure 5a, Figure S8a). Correlation with the additive relationship between parents was slightly lower (0.75) and non-significant at 5% (Figure S8a). No such strong correlation was found for per-cross PA in scenarios 1a or 1b (Figure S8a). The proportion of non-segregating markers showed low correlation with per-cross PA in all scenarios (Figure S8a).

**Figure 5:**
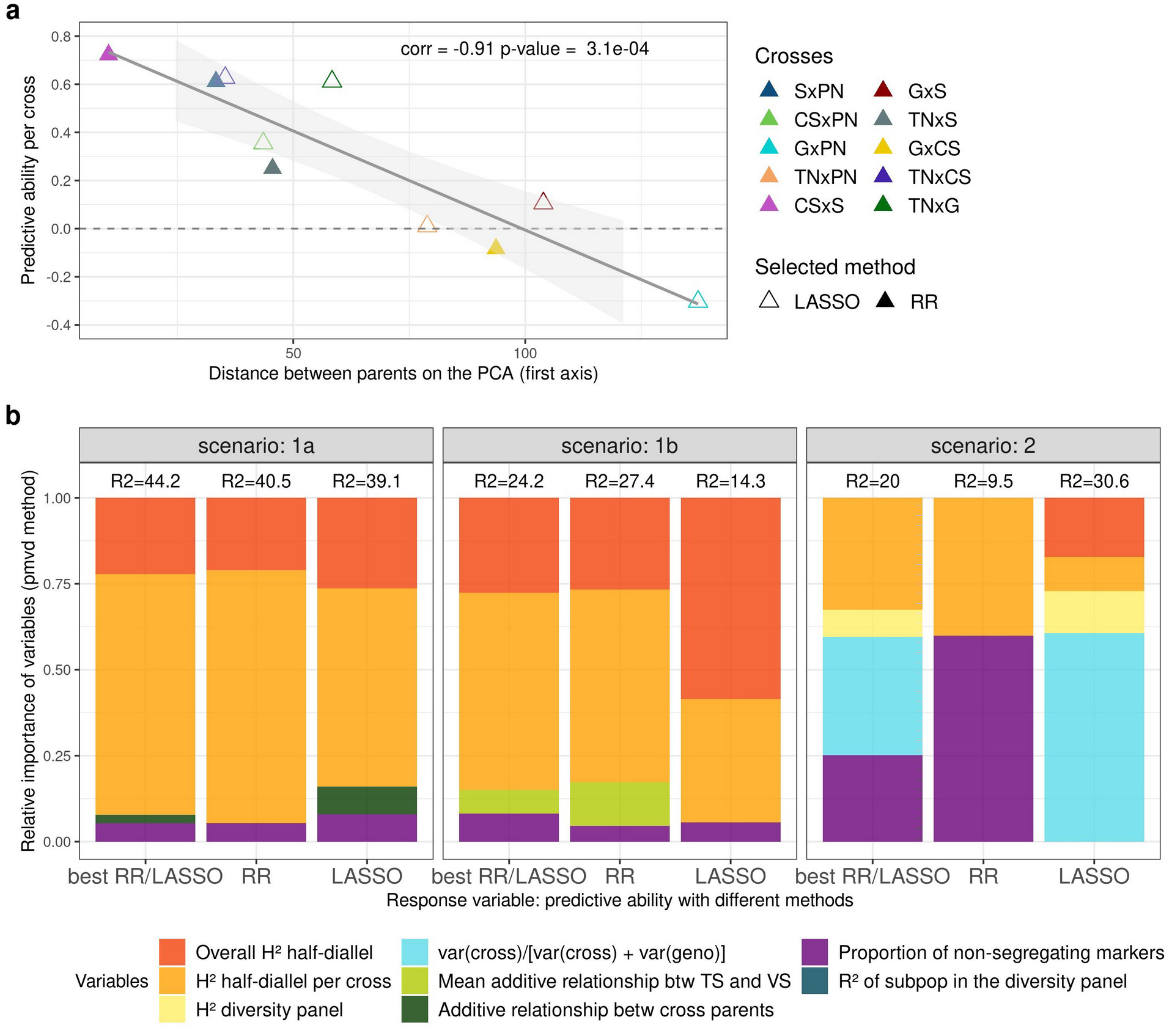
a: Plot of per-cross PA for cross mean in scenario 2, obtained with the best method between RR and LASSO for each cross, against the distance between cross parents on the first axis of the diversity panel PCA (Figure 1a). Best method is indicated with the triangle filling and cross with the color. b: Relative importance of variables affecting PA for Mendelian sampling in the three scenarios tested. Variables were selected from an overall model, after a model selection step. Response individual PA values were obtained either as the best one between RR and LASSO, with RR or with LASSO. Relative importance was estimated with pmvd method, from relaimpo R-package version 2.2-5.

Since variation in per-cross PA for scenario 2 was extremely large, from −0.3 for GxPN to 0.72 for CSxS (Figure 3a), we implemented TS optimization for each cross, to try and increase low PA values. Optimization actually improved PA for crosses with PA initially below 0.6, for TS sizes between 50 and 150 (Figure 6). The largest improvement, from −0.29 to 0.62, was observed for GxPN cross.

**Figure 6:**
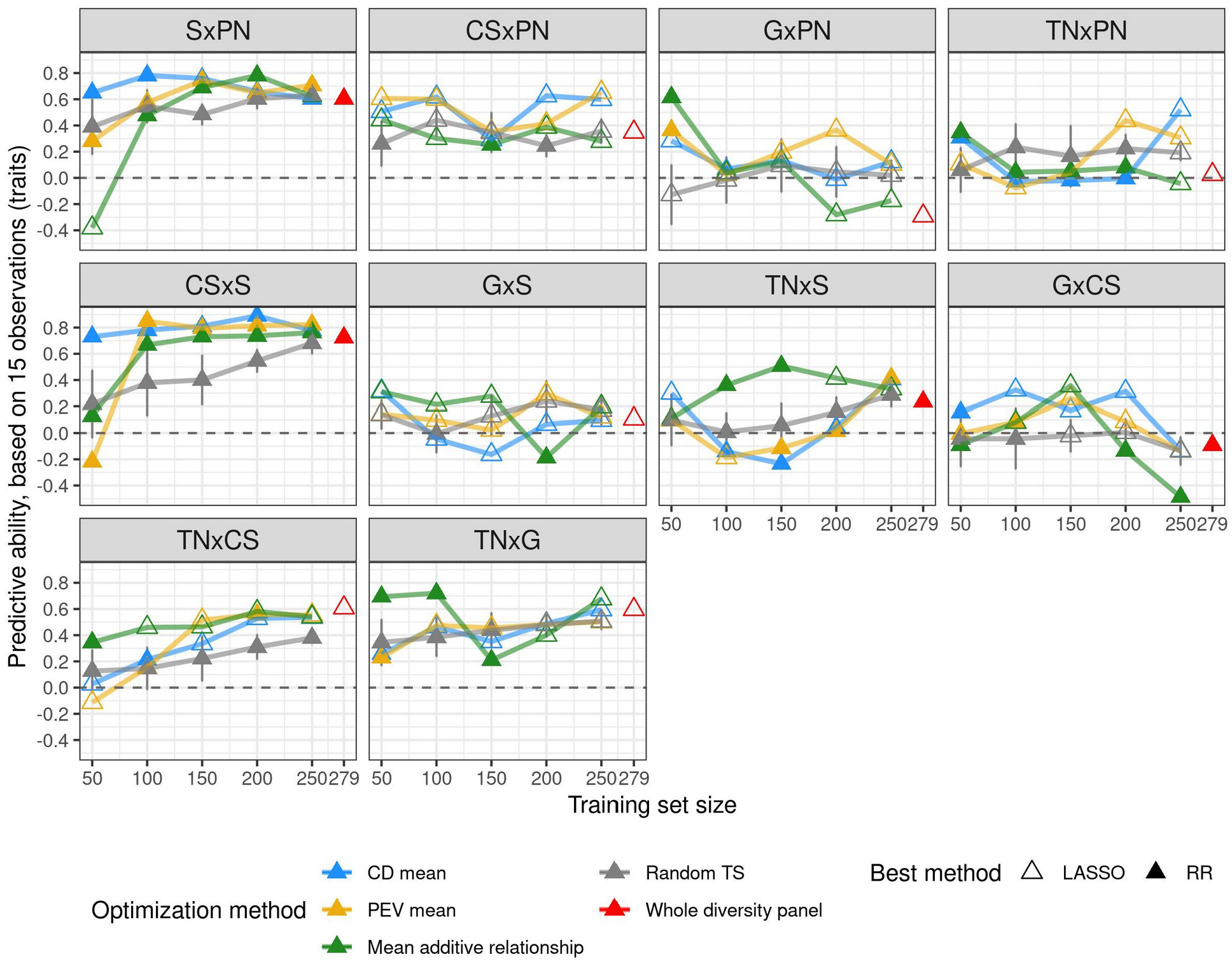
PA for cross mean predicion after training set optimization and with the best method between RR and LASSO, for each cross. Best method is indicated with the triangle filling and TS optimization method with the color. For comparison, random selection of TS genotypes (in grey) was performed and repeated ten times, error bars correspond to 95% of the confidence interval around the mean. We also report per-cross PA with the whole diversity panel (in red), with a maximum TS size of 279 which may vary depending on traits.

The variable that most affected per-trait PA was the 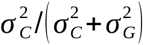 ratio (relative variance of cross effect). It was strongly correlated with PA in scenarios 1a and 1b (0.82 and 0.88, respectively), but not in scenario 2 (Figure S8b).

No other explanatory variable displayed any significant impact despite a fairly high correlation with per-trait or per-cross PA, which could be due to low sample sizes (15 and 10 for per-trait and per-cross PA, respectively).

#### Factors affecting Mendelian sampling prediction

To model Mendelian sampling PA for each scenario and method selected for each trait (RR, LASSO or best), we applied multiple linear regression on six to nine variables depending on the scenario, as detailed in Material and Methods. The highest coefficient of determination (44.2%) was obtained in scenario 1a with the best method (Figure 5b). Coefficients of determination were equivalent, lower and higher for LASSO compared to RR in scenarios 1a, 1b and 2, respectively. Three variables were found to impact PA in all scenarios: half-diallel overall H^2^, per-cross H^2^ and the proportion of non-segregating markers. Surprisingly, half-diallel overall H^2^ was not selected in scenario 2 with either RR or best method, while it had a strong effect in other modalities.

The selected variables were quite similar between scenarios 1a and 1b, with a high effect of half-diallel overall and per-cross H^2^, but differed in scenario 2 in which more variables were selected. Overall, most of the relative importance came from variables related to the trait and not to the genetic composition of TS or validation set (VS).

We also calculated individual PA with optimized TSs derived from the diversity panel (Figure S9). However, we did not observe any improvement compared to using the whole diversity panel. This is consistent with the fact that genetic relationship seemed not to impact PA (Figure 5b).

## Discussion

Our study allowed us to thoroughly explore GP potential in grapevine breeding, by scanning a large range of potentially useful configurations: (i) with 15 weakly related traits with variable levels of H^2^ and phenotypic structure (subpopulation or cross effects on phenotypic data) (Figure S4), (ii) in across-population scenarios with TS ranging from half-sibs (scenario 1b) to a diversity panel (scenario 2), (iii) with 10 balanced VS crosses. Moreover, we decomposed PA into cross mean and Mendelian sampling components, each being useful in breeding to select parental genotypes and offspring within crosses, respectively. All these results allowed us to get insight into main factors affecting PA. We will focus our discussion on prediction with the diversity panel as TS, since this is the most sought-after configuration in perennial species breeding.

### Range of PA values

For the prediction of cross mean, overall PA was 0.32 in scenario 2, equivalent to the average per-cross PA (0.29), while the average per-trait PA was twice as high (0.6) (Figure 3). In other studies concerning other plant crops, the average per-cross PA was not reported ^5,6,7,8^, probably because, in most cases, there were not enough traits to estimate it. Bernardo et al. ^5^ and Osthushenrich et al. ^6^ also reported a high-average per-trait PA, above 0.9, while Yamamoto et al. ^8^ reported PA values from 0.21 to 0.57 depending on the trait.

For the prediction of Mendelian sampling, overall average PA was slightly lower than overall PA for cross mean in scenario 2 (0.26 and 0.32, respectively). Yet, Mendelian sampling PA was still quite high, considering that TS was essentially unrelated to VS, i.e., with no first-degree relationship with predicted progenies. The same diversity panel was previously used in Flutre et al. ^26^ for predicting individual genotypic values of 23 additional Syrah x Grenache offspring. The reported PA for **mbw** was 0.56, whereas in the present study, we obtained 0.35 in the Grenache x Syrah progeny (n=59). We further investigated such discrepancy, and found it related to a sampling bias due to the small VS size in Flutre et al. ^26^ (data not shown).

The range of average per-trait Mendelian sampling PA observed in scenario 2 (from 0.15 to 0.38) was consistent with those described on fruit perennial species where individual prediction was performed with a TS not directly related to the VS (neither half-sib nor full-sib). In *Coffea*, Ferrao et al. ^28^ reported differences in per-trait PA, from slightly negative values up to ca. 0.60. But, in this study, overall PA was calculated for all crosses of the VS, thus encompassing both cross mean and Mendelian sampling predictions, making comparison with our Mendelian sampling results alone impossible. In contrast, some studies in apple yielded within cross individual PA values. For instance, Muranty et al. ^29^ reported average per-trait PA ranging from −0.14 to 0.37, and Roth et al. ^30^ found PA values from −0.29 to 0.72 for fruit texture, highly dependent on the cross for all traits. Conversely, our PA values were mainly stable over crosses and variable over traits, in the three scenarios (Figure S7). This difference might partly be due to the larger trait diversity we explored as compared to Roth et al. ^30^, as suggested by comparing our Figure S4 with their Figure 1A. A complementary explanation could be that progeny size varied from 15 to 80 in Roth et al. ^30^, while here progeny sizes were very close and thus less likely prone to sampling variability and to upward or downward bias.

Several factors may influence Mendelian sampling PA in our study compared to others. Among potential inflating factors, we can mention a slight over-representation of phenotyped individuals from the WW panel subpopulation, to which four out of the five parents of the half-diallel belong, leading to a higher genetic relationship between effective TS and VS. Factors potentially decreasing PA could be differences between TS and VS experimental designs since the diversity panel and the half-diallel were not phenotyped on the same years, had different plant management systems (overgrafting or simple grafting, respectively) and were planted a few kilometers apart. Nevertheless, for most studied traits, two years of phenotyping were used to compute genotypic BLUPs, which could at least compensate for differences between years, usually referred to as the millesime effect.

### Variables affecting PA in across-population genomic prediction

We focused on PA obtained with the best method between RR and LASSO, to take into account the part of variability among traits associated with genetic architecture. Indeed, LASSO is supposed to be better adapted to traits underlined by few QTLs, while RR would yield better PA for highly polygenic traits. However, we showed that for a given trait x cross combination, i.e., for a given genetic architecture, the best method selected changed depending on the scenario: LASSO was more often selected for scenario 2 than for scenario 1a, both for cross mean and individual prediction. This means that the best method choice also depends on the relationship between TS and VS. This was also suggested in cattle breeding by MacLeod et al. ^31^, who found that BayesRC method (comparable to LASSO) yielded better results than GBLUP (comparable to RR) for across-population GP.

Regarding the other factors affecting PA, for cross mean prediction in scenario 2, no tested variable significantly affected per-trait PA. Conversely, per-cross PA was strongly affected by the genetic distance between parents (Figure 5a, Figure S8a). To our knowledge, such correlation has never been reported before, most probably because previous works investigated too few traits to afford per-cross PA calculation. We could hypothesize that when one parent is farthest from WW -the most represented panel subpopulation in TS- (e.g., Grenache, Figure 1a, Figure S1), marker effects for this parent might reflect different QTLs or allelic frequencies, compared to WW ones, thereby explaining the decrease in PA for crosses related to Grenache. Such differences underlying marker effects were already described in maize ^32^. Simultaneously, some QTLs in this parent might be less genetically linked to causal polymorphisms due to more recombinations. However, this cannot be the only explanation for the large correlation of per-cross PA with pairwise parent distance, because the correlation between PA and genetic distance between TS and VS was much lower (Figure S8a).

For the prediction of Mendelian sampling, the variables explaining individual PA in scenario 2 were quite different from those explaining cross mean PA. Trait-related variables had a large impact on individual PA: half-diallel overall and per-cross heritability, but also the relative variance of cross effect (Figure 5b). Surprisingly, genetic relationship between TS and VS had little to no impact on PA, although this factor has often been reported to affect PA ^17,15^. Most studies reported separately the effects of different variables on individual PA. Riedelsheimer et al. ^33^ also performed multiple linear regression of individual PA on several factors to study their impact. They found that TS composition (number of crosses and their relationship with VS) explained most of the variance (41.7 %), followed by trait (27.6%) and VS composition (4.8%). The variance in genetic relationship between TS and VS may be smaller in our study.

### Practical consequences on breeding programs

Across-population GP with model training in a diversity panel appeared to be promising in grapevine breeding for some traits and crosses, particularly for parent choice (Figure 3; Figure 4; Table S3; Figure S7).

The usefulness of GP for better selecting parents for future crosses can be at first assessed by the low overall correlation between mean parental genotypic values (BLUPs) and mean offspring BLUPs (0.28; see also Figure S10). This correlation was much lower than overall PA for cross mean in scenario 1b (0.66) and slightly lower than overall PA for cross mean in scenario 2 (0.32). In strawberry, Yamamoto et al. ^8^ also evidenced the interest of GP for predicting cross mean, with no additional benefit from including dominance effects into GP models, even if cross means were not equal to parental means. Moreover, in some cases, GP could provide other advantages over mean parental genetic values, for instance when parents are not phenotyped for some reasons, because too young or without representative phenotypes (e.g., using microvine ^34^, in a new environment, etc). This was actually the case, in our half-diallel trial, for the Terret Noir parent, which suffered from mortality probably due to rootstock incompatibility and consequently had no phenotypic record for most studied traits.

Even though PA was quite high for some traits and crosses in scenario 2, on average it remained moderate both for cross mean and individual prediction. Both PAs were much higher in scenario 1a, due to increased relationship between training and validation sets. Nevertheless, such an extreme configuration is rarely used in plant breeding programs, especially in perennial species, because it requires to partly phenotype the cross to be predicted. An intermediate configuration, scenario 1b, could be implemented in breeding programs when PA from scenario 2 is not sufficient and half-sib families are available, because in this scenario, cross mean PA was similar as in scenario 1a and individual PA intermediate between scenarios 1a and 2.

We found TS optimization useful mostly for cross mean prediction for crosses with low PA. The advantage of TS optimization was less clear for individual prediction. This was consistent with the fact that genetic parameters more strongly affected cross mean PA than individual PA. In contrast, Roth et al. ^30^ observed in apple a systematic increase of individual PA with an optimized TS in the same context (i.e., with a diversity panel as TS and bi-parental families as VS, and common optimization methods). To our knowledge, only a single study tested TS optimization for cross mean prediction, by Heslot and Feoktistov ^35^, who implemented optimization of parent selection for hybrid crossing in sunflower while selecting individuals to phenotype, but did not calculate cross mean PA.

Since our results show that prediction of cross mean can be quite accurate and useful in scenario 2, we decided to go one step further and implemented cross mean prediction for all 38,781 possible crosses between the 279 genotypes of the diversity panel, based on parental average genotypes (Table S2) and on marker effects estimated with RR in this population. As predicted cross mean were biased for some traits in the ten half-diallel crosses (Figure S5), we estimated the bias for each trait from these data to correct the predicted mean in the possible diversity panel crosses. Figure S11 shows the large potential diversity to be explored through crossing in grape, for all the traits considered in the present study, illustrating the finding of Myles et al. ^36^ that genetic diversity in grapevine was largely unexploited. Such an example opens many prospects for the use of GP to design future crosses. Indeed, we limited here our prediction to the 279 panel genotypes representing the *Vitis vinifera* diversity, but potentially any other (unphenotyped) genotype of interest with dense genotypic data could be used for this purpose as exemplified with the half-diallel, since its five parents were not part of the diversity panel.

### Prospects

Based on our results, the following improvements could be tested: i) increase SNP density ^25, 37^ and include structural variants ii) implement non-additive effects in GP models such as dominance or epistatic effects and iii) add crosses from other panel subpopulations as VSs. Indeed, since all our half-diallel crosses had at least one parent belonging to the WW subpopulation, it would be beneficial to include crosses with parents from the WE and TE subpopulations too. Specific GP models that include genetic structure in marker effect estimation ^38, 39^ could also be tested.

Predicting cross variance could also prove useful to design the offspring selection step, more specifically for choosing the number of offspring to test or produce for a given cross. Depending on the available funds and breeding program, a breeder may want to select crosses with high genetic variance, in order to maximize the probability to generate top-ranking genotypes. Conversely, choosing a cross with low variance could limit the risk of breeding poor genotypes.

### Conclusion

We implemented GP in grapevine in a breeding context, i.e., across populations, on 15 traits, in ten related crosses, and obtained moderate to high PA values for some crosses and traits, thus showing GP usefulness in grapevine. Never before had genomic prediction been implemented for so many traits and crosses simultaneously in this species. We showed that per-cross PA was strongly correlated with the genetic distance between parents, whereas Mendelian sampling PA was largely determined by trait-related variables, such as heritability and the magnitude of the cross effect.

## Material and Methods

### Plant material

The half-diallel consists of 10 pseudo-*F*_1_ bi-parental families obtained by crossing five *Vitis vinifera* cultivars: Cabernet-Sauvignon (CS), Pinot Noir (PN), Terret Noir (TN), Grenache (G) and Syrah (S) ^40^. Each family comprised between 64 and 70 offspring, with a total of 676 individuals including parents.

The diversity panel consists of 279 cultivars selected as maximizing genetic diversity and minimizing kinship among cultivated grapevine. Grapevine genetic diversity is highly heterozygous and weakly structured into three subpopulations: WW (Wine West), WE (Wine East) and TE (Table East) ^27^.

### Field experiments

#### Field design

The half-diallel was created in 1998 at INRAE Montpellier, grafted on Richter 110, and planted in 2005, at the Institut Agro experimental vineyard “Le Chapitre” in Villeneuve-lès-Maguelone (Southern France). The progenies were planted in two randomized complete blocks, with plots of two consecutive plants per offspring per block.

The field design for the diversity panel was previously described in Flutre et al. ^26^. Briefly, cultivars were overgrafted on 6-year-old Marselan in 2009, itself originally grafted on Fercal rootstock, a few kilometers away from the diversity panel. They were planted in five randomized complete blocks, with one plant per cultivar per block.

#### Phenotyping

We studied 15 traits in both trials: berry composition with malic (**mal.ripe**), tartaric (**tar.ripe**) and shikimic acid (**shik.ripe**) concentrations in *μ_eq_* · *L*^−1^ measured at ripe stage (20° Brix) (according to Rienth et al. ^41^), from which two ratios were derived, shikimic / tartaric acid (**shiktar.ripe**) and malic / tartaric acid (**maltar.ripe**); morphological traits with mean berry weight (**mbw**, in g) measured on 100 random berries, mean cluster weight (**mcw**, in g), mean cluster length (**mcl**, in cm) and mean cluster width (**mcwi**, in cm), measured on 3 clusters, number of clusters (**nbclu**) and cluster compactness (**clucomp**) measured on the OIV semi-quantitative scale; phenology traits with veraison date (onset of ripening; **verday**, in days since January 1st), maturity date corresponding to berries reaching 20° Brix (**samplday**, in days since January 1st) and the interval between veraison and maturity (**vermatu**, in days); vigour (**vigour**, in kg), derived as the ratio between pruning weight and the number of canes. Phenotypic data were collected between 2013 and 2017 for the half-diallel and in 2011-2012 for the diversity panel. There was a slight over-representation of phenotypes from the WW subpopulation because of fertility issues in WE and TE subpopulations

### SNP genotyping

For the half-diallel, we used genotyping-by-sequencing (GBS) SNP markers derived by Tello et al. ^40^, 622 of the 676 individuals being successfully genotyped, as well as the five parents. Raw GBS data were processed separately for each cross, and then markers from all crosses were merged together (390,722 SNPs), thus generating many missing data (85% of missing data per marker on average), since all markers did not segregate in all progenies. Markers with more than 80% of missing data were removed and remaining markers were imputed with FImpute3 ^42^ (86,017 SNPs). Some parental cultivars were used either as female or male, depending on the cross, a configuration not allowed by FImpute3. We thus declared only a partial pedigree maximizing the number of crosses defined with both parents (Table S4). For the diversity panel, we used the same SNP markers as in Flutre et al. ^26^, except that we applied a filter on minor allelic frequency (5%) and no filter on linkage disequilibrium, which yielded 83,264 SNPs.

Finally, we only retained the 32,894 SNPs common to both populations.

### Phenotypic data analyses

#### Half-diallel

##### Statistical modeling for estimating genotypic values

For each trait, we excluded outlier values by visual inspection of raw phenotypic data and computed a log or square-root transformation if its distribution looked skewed. Then, we fitted the following linear mixed full model by Maximum Likelihood:

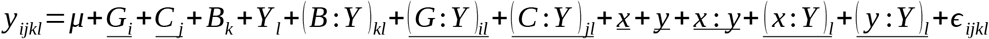

with *y_ijkl_* the phenotype of genotype i from cross j in block k and year l. Among the fixed terms, *μ* was the overall mean, and *B_k_* and *Y_l_* the effects of block k and year l. Among the random terms, *G_i_* and *C_j_* were the effects of genotype i nested within cross j, and *x* and *y* the field coordinates. Interactions are indicated with “:”. *ε_ijkl_* was the random residual term, assumed to be normally distributed.

Sub-model selection was based on Fisher tests for fixed effects and log-likelihood ratio tests for random effects. It was performed with the step function from lmerTest R-package ^43^. Variance components were estimated after re-fitting the selected model by Restricted Maximum Likelihood, and diagnostic plots were drawn to visually check the acceptability of model hypotheses such as homoscedasticity or normality. Best Linear Unbiased Predictors (BLUPs) of cross (C) and genotype (G) values were computed. For genomic predictions, we used their sum (C+G) as total genotypic values for both training and validation data. Variance component estimates were used to compute the proportion of genetic variance due to differences between crosses as: 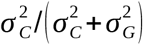.

##### Heritability estimation

We estimated overall (for the whole half-diallel) broad-sense heritability for genotype-entry means ^44^ as:

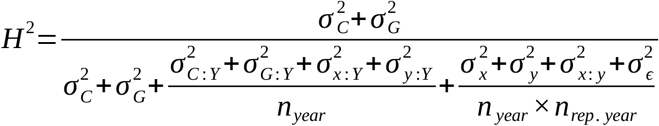

with genotype (G) and cross (C) variances at the numerator. Random variance components involving year (Y) were divided by the mean number of years (*n_year_*). Other random variance components involving spatial effects or residuals were divided by the mean number of years times the mean number of replicates per year (*n_rep · year_*).

We also estimated broad-sense heritability per cross (thereafter used to name half-diallel full-sib family). For that, we applied the same selected model, but removed all effects involving cross. Then, we estimated variance components within each cross, and heritability with the same formula, after removing variances involving cross.

All information on analyses of phenotypic data and heritability of the half-diallel is detailed in Table S1.

#### Diversity panel

We used the genotypic values previously estimated in Flutre et al. ^26^ with a similar statistical procedure to the one described above for the half-diallel. All phenotypic analysis information is provided in Table S3 of Flutre et al. ^26^.

For each of the two populations, genotypic BLUPs were scaled, allowing comparison among traits.

#### Genomic prediction statistical methods

Marker effects were estimated using two methods to take into account varying genetic architecture among the traits studied. Ridge regression (RR) ^45^, best adapted to many minor QTLs, shrinks marker effects towards 0. Least Absolute Shrinkage and Selection Operator (LASSO) ^46^, best adapted to a few major QTLs, applies a L1 norm on allelic effects, thus forcing some to be exactly 0. Both methods were implemented with R/glmnet package ^47^ and the amount of shrinkage, controlled by *λ* parameter, was calibrated by five-fold inner cross-validation within each training set, using cv.glmnet function.

#### Genomic prediction scenarios

We assessed prediction within half-diallel crosses under three different training scenarios (Figure 2; Table S5):

- **Scenario 1a:** whole half-diallel prediction. We applied random outer 10-fold cross-validation over the whole half-diallel population. In each fold, 90% of the phenotyped offspring were used as the training set (TS) and the remaining 10% as the validation set (VS). Cross-validation was replicated ten times.
- **Scenario 1b**: half-sib prediction. For each half-diallel cross used as VS, we trained the model with the three half-sib crosses of each parent in turn, thus predicting each cross twice.
- **Scenario 2**: across-population prediction. We used the whole diversity panel as TS and each half-diallel cross as VS.

#### Predictive ability assessment

In order to account for the effect of genetic architecture, we applied both RR and LASSO methods for each trait and cross and kept the best PA, for both cross mean and within cross individual prediction.

##### Prediction of cross mean

Cross mean PA was assessed as Pearson’s correlation between the average value of observed total genotypic values (sum of genotype and cross BLUPs for each offspring) for each cross, and the mean predicted genotypic value per cross, calculated in two ways, as:

- average predicted value over all offspring of the cross. In scenario 1a, each offspring was predicted 10 times, thus we also averaged the predicted value over the 10 replicates.
- predicted value for the parental average genotype, defined at each locus and for each cross as the mean allelic dosage according to the expected segregation pattern based on parents’ genotypes (Table S2).
- genotypic values predicted with these two modalities were highly correlated (above 0.98) in the three scenarios and for the two methods (partly shown in Figure S12). Therefore, in subsequent analyses, we used only prediction with parental average genotypes.

Pearson’s correlation between observed and predicted values was calculated on all cross x trait combinations (overall PA), for each trait (per-trait PA) and for each cross (per-cross PA).

##### Within-cross individual prediction

We measured PA within each cross in each scenario as Pearson’s correlation between observed total genotypic values and predicted genotypic values.

#### Test of variables affecting predictive ability

We tested the effect of several variables on within-cross individual PA, in each scenario. We built a multiple linear regression model with PA per trait x cross combination as the response variable and as predictors, a set of variables common to all three scenarios plus specific variables for scenarios 1b and 2. Common variables were: the proportion of non-segregating markers in the cross, overall and per-cross broad-sense heritability, the distance between the parents of the cross measured either as the additive relationship or as the distance on the first or first two axes of the panel PCA (Figure 1a) and the proportion of genetic variance due to differences between crosses (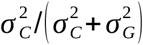 ratio). A specific variable for scenarios 1b and 2 was the mean additive relationship between training and validation sets. In scenario 2, it was calculated for each trait only with phenotyped individuals. Specific variables for scenario 2 were: broad-sense heritability in the diversity panel (retrieved from Flutre et al. ^26^ and Table S1) and the percentage of trait variance explained by the subpopulation factor (see below). After fitting the overall model, we applied a forward-backward stepwise regression, with the AIC criterion to select the best explanatory model. Then, we estimated the relative importance of each variable selected in this model with the pmvd method ^48^, which allows to decompose the R^2^ of correlated regressors with the R-package relaimpo ^49^.

The percentage of trait variance within the diversity panel explained by subpopulation (WW, WE or TE) was evaluated by fitting for each trait the following linear model: *G* = *P* + *ε*, where *G* is the genotypic (BLUP) value within the diversity panel, *P* is a fixed subpopulation effect, and *ε* a random residual term. The percentage of variance due to differences between subpopulations was then estimated as the coefficient of determination (R^2^) of the model.

#### Training set optimization

We tested three methods for optimizing TS in scenario 2, for both cross mean and within-cross individual prediction. We used the STPGA R-package^50^ to implement Prediction Error Variance (*PEVmean*) and *CDmean* (based on the coefficient of determination)^10^. Moreover, we computed the mean relationship criterion (*MeanRel*), as the mean additive relationship between each genotype in TS and all genotypes in VS. Each optimized TS was specific to a cross. The realized additive relationship based on marker data was estimated using the rrBLUP R-package^51^ with the A.mat function implementing the formula from VanRaden et al. ^52^. For each of these three optimization methods, we tested five TS sizes (50, 100, 150, 200, 250). PA values obtained with each optimized TS were compared with those obtained with a random sample of genotypes of the same size, repeated 10 times.

#### Supporting information

Supplementary tables and figures

## Data availability

All analyses were conducted using free and open-source software, mostly R. Phenotypic and genotypic data, R scripts and result tables are available at https://data.inrae.fr/privateurl.xhtml?token=1925c973-a11b-45ad-b297-69db8ec2c270.

## Acknowledgments

We thank Charles Romieu for his help on the acquisition of phenotypic data as well as Philippe Châtelet and Morgane Roth for their suggestions on the text. This work has been realized with the support of the SouthGreen platform and MESO@LR-Platform at the University of Montpellier.

## Author contribution statement

VS, AD, PT, TF, JPP and LLC conceived the idea of the study and contributed to funding acquisition; TP, PF, AD and JPP obtained the phenotypic data used in this work; VS, AD, PT, LLC, TF and CB performed and interpreted results; AD conceived the half-diallel population; PT is the PhD supervisor of CB; CB wrote the original draft, which was reviewed and edited by all authors. All authors read and approved the final manuscript.

