## Supplementary tables and figures for "Across-population genomic prediction in grapevine opens up promising prospects for breeding"

#### Table of contents

Figure S1 Additive relationship between half-diallel parents and diversity panel cultivars. 2

Table S1 Information on mixed model selection and genotypic BLUP estimation for 15 traits in the half-diallel population 3

Figure S2 Per cross broad-sense heritability in the half-diallel. 4

Figure S3 Distribution of genotypic value estimates (BLUPs) for 15 traits, in each diversity panel subpopulation and each half-diallel cross 9

Figure S4 PCA applied to genotypic BLUPs for the 15 traits. 10

Table S2 Computation of parental average genotypes. 11

Table S3 Predictive ability of cross mean. 12

Figure S5 Observed vs predicted cross means for each trait in the half-diallel 12

Figure S6 Observed vs predicted individual genotypic values for 15 traits. 20

Figure S7 Distribution of predictive ability for Mendelian sampling genomic prediction. 21

Figure S8 Correlation plot for PA of cross mean and potential explanatory variables. 22

Figure S9 Distribution of predictive ability for Mendelian sampling genomic prediction, after training set optimization 23

Figure S10 Mean offspring observed genotypic value vs parental average observed genotypic value in each half-diallel cross for 15 traits 23

Figure S11 PCA of predicted cross mean genotypic values for all 38,781 possible simulated crosses between the 279 varieties of the diversity panel 24

Table S4 Partial pedigree of half-diallel crosses used for marker imputation. 24

Table S5 Training and validation sets composition for each scenario used to assess genomic prediction 25

Figure S12 Observed vs predicted cross mean for 15 traits. 25

Bibliography 25

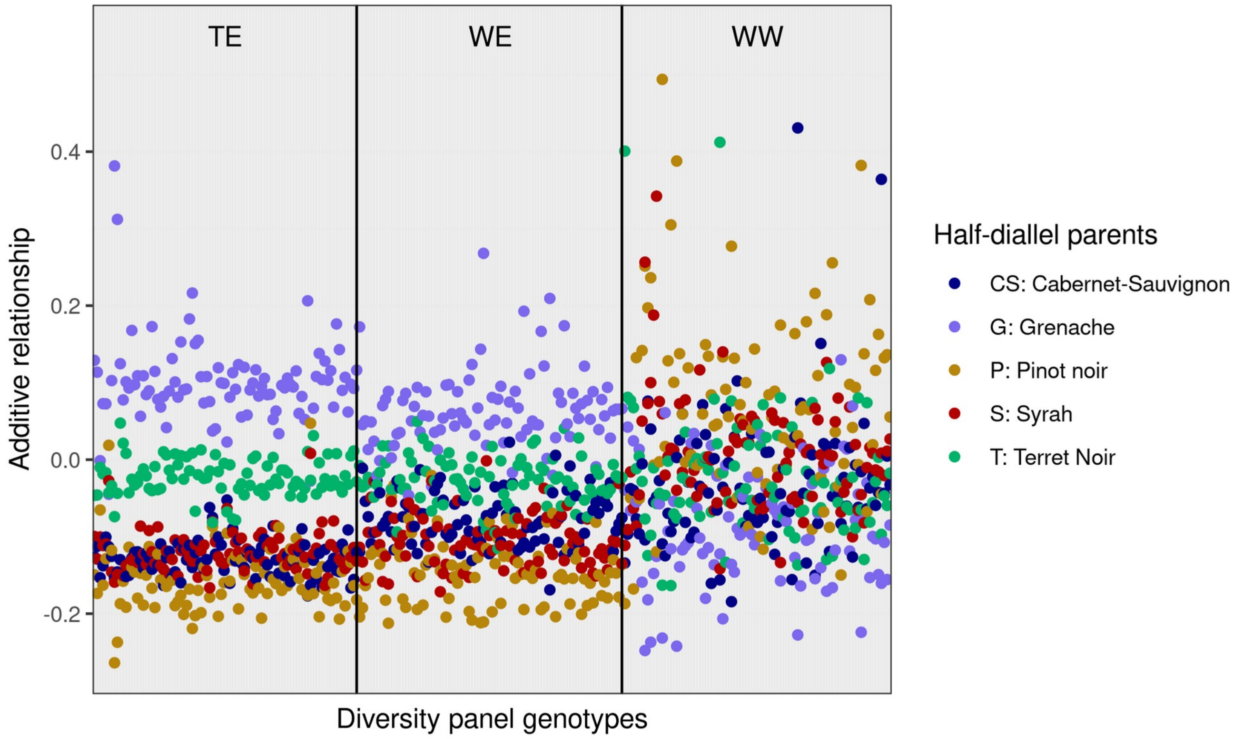


#### Figure S1 Additive relationship between half-diallel parents and diversity panel cultivars.

Additive relationship was calculated with the VanRaden (2008) method, with 32,894 SNPs. For each diversity panel genotype, its additive relationship with the five half-diallel parents was computed. Each point corresponds to one cultivar in the diversity panel, ordered by on the *x*-axis subpopulation (TE: Table East, WE: Wine East, WW: Wine West), and colored according to the corresponding half-diallel parent genotype.

| trait | years | missing.data.percent | transformation | fixed.effects | random.effects |
|---|---|---|---|---|---|
| **mal.ripe** | 2015 | 38 | NA | none | geno, cross, x |
| **tar.ripe** | 2015 | 38 | NA | none | geno, cross, x |
| **shik.ripe** | 2015 | 42 | log | block | geno, cross, x |
| **shiktar.ripe** | 2015 | 42 | log | block | geno, cross, x |
| **maltar.ripe** | 2015 | 38 | log | none | geno, cross, x |
| **verday** | 2013, 2014, 2017 | 36 | NA | block, year, block:year | geno, cross, x, geno:year, cross:year |
| **sampldday** | 2013, 2014, 2015 | 28 | NA | block, year | geno, cross, geno:year, year:x |
| **vermatu** | 2013 | 33 | NA | block | geno, cross, x |
| **clucomp** | 2013, 2014, 2015 | 30 | NA | block, year, block:year | geno, cross, geno:year, cross:year, year:x |
| **nbclu** | 2013, 2014, 2015 | 28 | sqrt | block, year, block:year | geno, cross, x:y, geno:year, cross:year, year:x |
| **mcl** | 2013, 2014, | 28 | NA | year | geno, cross, x:y, geno:year, cross:year, |

| | | | | | |
|---|---|---|---|---|---|
| | 2015 | | | | year:x, year:y |
| **mcwi** | 2013 | 32 | NA | block | geno, cross |
| **mcw** | 2013, 2014, 2015 | 29 | log | block, year, block:year | geno, cross, x:y, geno:year, cross:year, year:x |
| **mbw** | 2013, 2014, 2015 | 29 | sqrt | block, year, block:year | geno, cross, x:y, geno:year, cross:year |
| **vigour** | 2014, 2015 | 16 | log | block, year, block:year | geno, cross, x:y, geno:year, cross:year, year:x, year:y |

| trait | var.geno | var.cross | H2 | H2.low | H2.high | CV.geno | CV.geno.low | CV.geno.high | var.cross.geno |
|---|---|---|---|---|---|---|---|---|---|
| **mal.ripe** | 460 | 85.94 | 0.58 | 0.55 | 0.61 | 0.17 | 0.14 | 0.19 | 0.16 |
| **tar.ripe** | 230 | 50.68 | 0.70 | 0.68 | 0.73 | 0.16 | 0.14 | 0.18 | 0.18 |
| **shik.ripe** | 0.379 | 0.40 | 0.80 | 0.78 | 0.82 | 0.26 | 0.22 | 0.31 | 0.51 |
| **shiktar.ripe** | 0.377 | 0.43 | 0.84 | 0.82 | 0.86 | 0.09 | 0.08 | 0.10 | 0.53 |
| **maltar.ripe** | 0.042 | 0.02 | 0.74 | 0.71 | 0.76 | 0.73 | 0.55 | 1.00 | 0.31 |
| **verday** | 13.1 | 3.84 | 0.8 | 0.79 | 0.81 | 0.02 | 0.01 | 0.02 | 0.23 |
| **samplday** | 36.2 | 30.63 | 0.82 | 0.81 | 0.84 | 0.02 | 0.02 | 0.03 | 0.46 |
| **vermatu** | 48.0 | 19.30 | 0.65 | 0.63 | 0.68 | 0.18 | 0.15 | 0.20 | 0.29 |
| **clucomp** | 1.53 | 0.16 | 0.81 | 0.80 | 0.83 | 0.23 | 0.21 | 0.24 | 0.09 |
| **nbclu** | 0.742 | 0.15 | 0.84 | 0.83 | 0.85 | 0.20 | 0.18 | 0.22 | 0.17 |
| **mcl** | 2.25 | 0.53 | 0.80 | 0.78 | 0.81 | 0.12 | 0.11 | 0.13 | 0.19 |
| **mcwi** | 1.05 | 0.58 | 0.50 | 0.46 | 0.52 | 0.12 | 0.10 | 0.15 | 0.35 |
| **mcw** | 0.083 | 0.05 | 0.83 | 0.82 | 0.84 | 0.05 | 0.05 | 0.06 | 0.36 |
| **mbw** | 0.019 | 0.02 | 0.92 | 0.91 | 0.93 | 0.1 | 0.09 | 0.11 | 0.51 |
| **vigour** | 0.096 | 0.01 | 0.77 | 0.76 | 0.80 | 0.12 | 0.11 | 0.13 | 0.06 |

#### Table S1 Information on mixed model selection and genotypic BLUP estimation for 15 traits in the half-diallel population.

**trait:** see abbreviations meaning in Methods section.

**years:** years in which the trait was phenotyped.

**missing.data.percent**: percentage of missing raw phenotypic data, relative to the full initial design.

**transformation**: transformation applied to raw phenotypic data, before model selection and BLUP estimation (NA: none; sqrt: square root; log: neperian logarithm).

**fixed.effects / random.effects**: fixed and random effects kept in the final selected model, as defined in Methods section.

**var.geno**: intra-cross genotypic variance estimate (534 to 624 levels, depending on the trait).

**var.cross**: cross variance estimate (10 levels)

**H2 / H2.low / H2.high:** broad-sense heritability estimate and its confidence interval bounds computed through bootstrapping.

**CV.geno / CV.geno.low / CV.geno.high**: estimated coefficient of variation of genotypic effect and its confidence interval bounds.

**var.Cross.geno**: $\sigma_C^2/\left(\sigma_C^2+\sigma_G^2\right)$ ratio, as defined in Methods section.

### Figure S2 Per cross broad-sense heritability in the half-diallel.

For crosses and traits, see abbreviation meaning in Methods section.

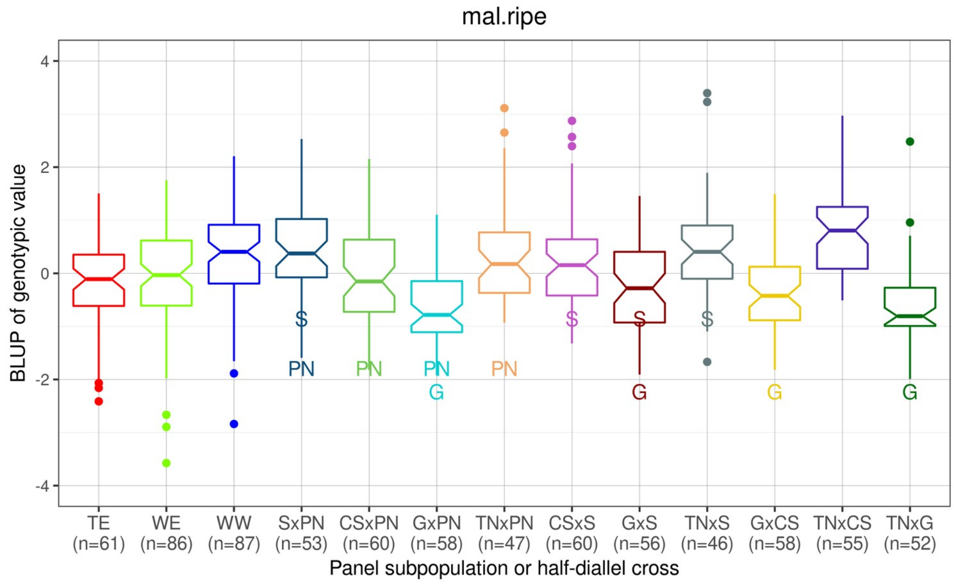

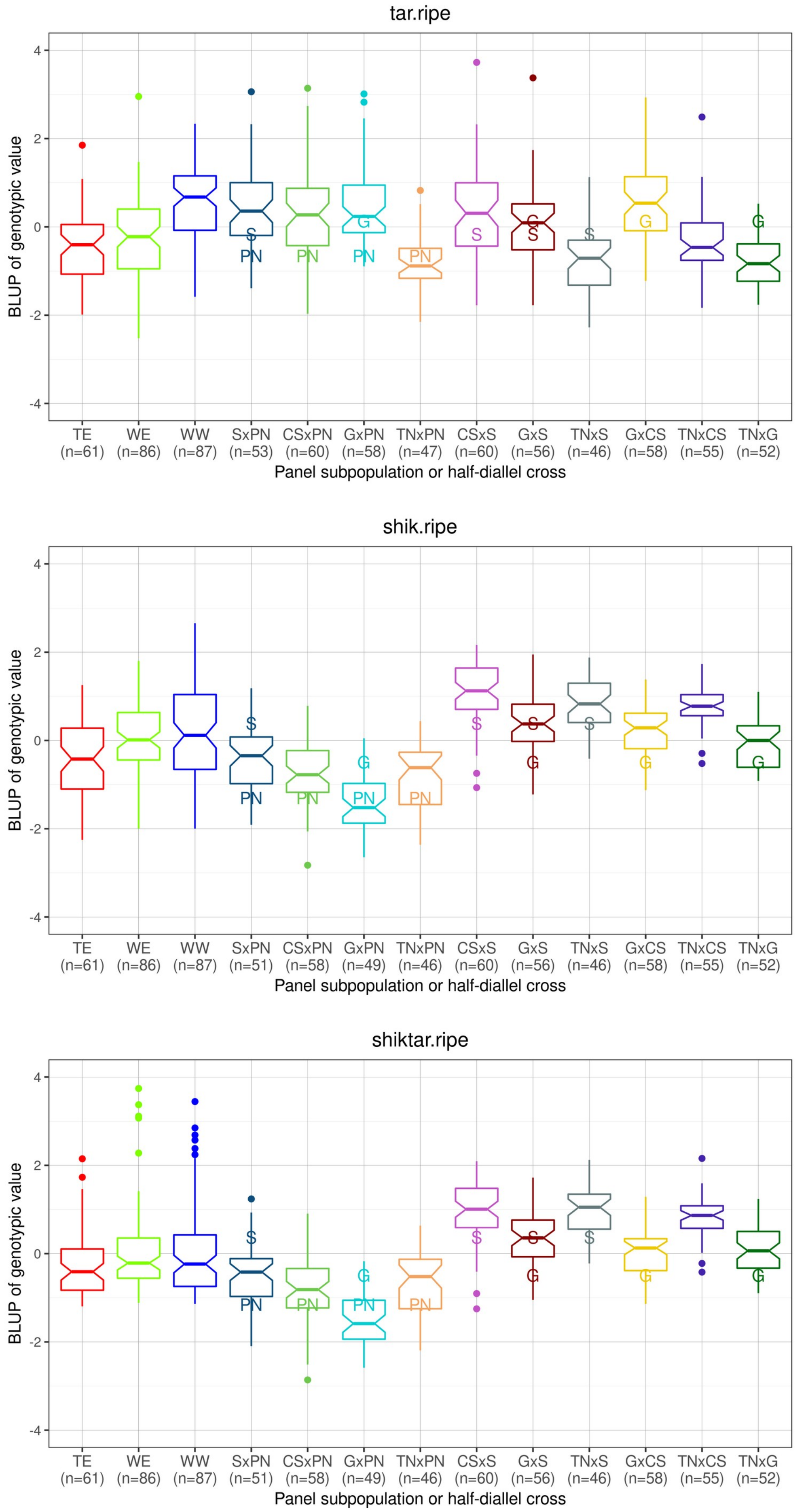

tar.ripe
BLUP of genotypic value
TE (n=61) WE (n=86) WW (n=87) SxPN (n=53) CSxPN (n=60) GxPN (n=58) TNxPN (n=47) CSxS (n=60) GxS (n=56) TNxS (n=46) GxCS (n=58) TNxCS (n=55) TNxG (n=52)
Panel subpopulation or half-diallel cross
shik.ripe
BLUP of genotypic value
TE (n=61) WE (n=86) WW (n=87) SxPN (n=51) CSxPN (n=58) GxPN (n=49) TNxPN (n=46) CSxS (n=60) GxS (n=56) TNxS (n=46) GxCS (n=58) TNxCS (n=55) TNxG (n=52)
Panel subpopulation or half-diallel cross
shiktar.ripe
BLUP of genotypic value
TE (n=61) WE (n=86) WW (n=87) SxPN (n=51) CSxPN (n=58) GxPN (n=49) TNxPN (n=46) CSxS (n=60) GxS (n=56) TNxS (n=46) GxCS (n=58) TNxCS (n=55) TNxG (n=52)
Panel subpopulation or half-diallel cross

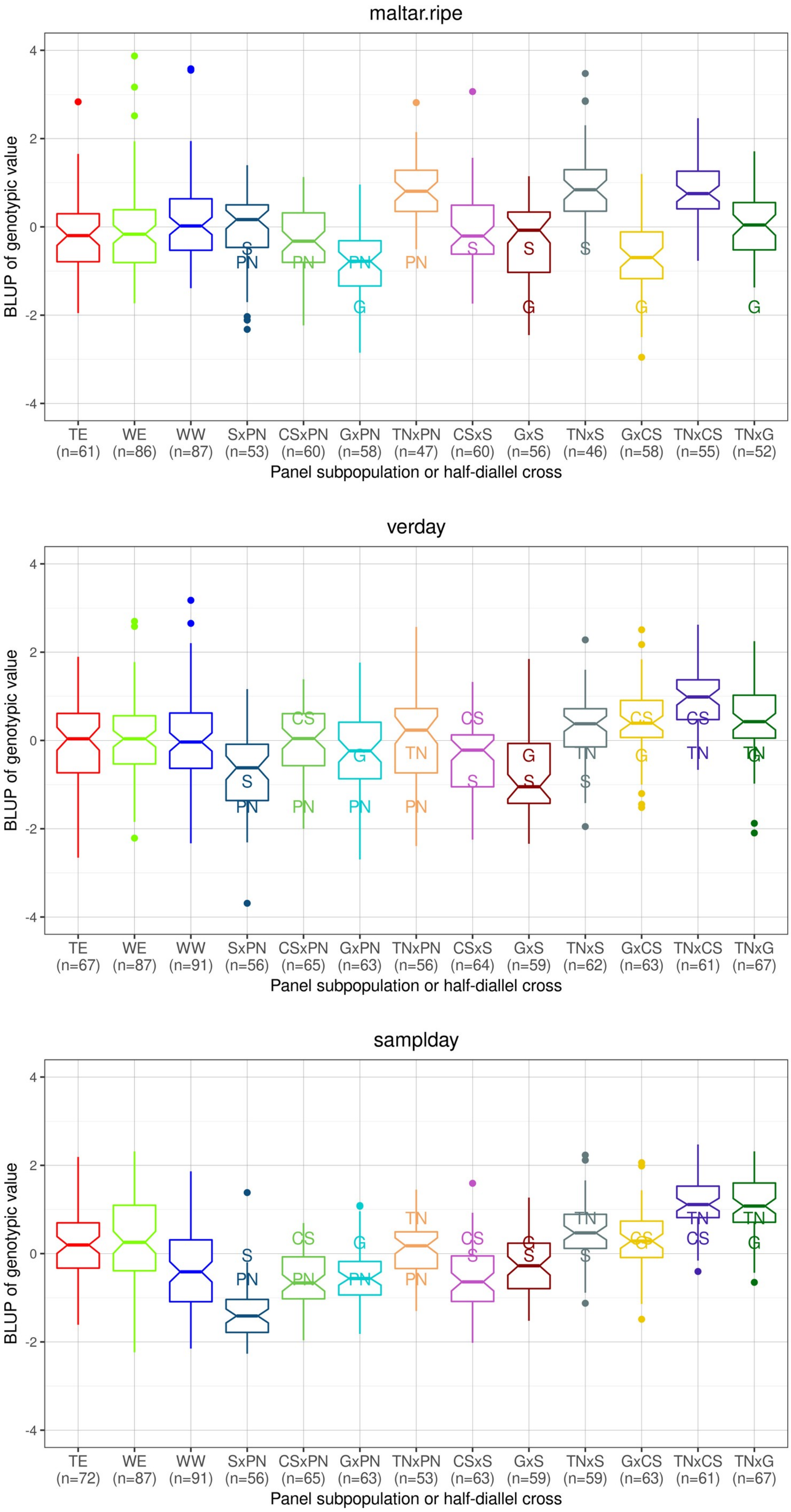

maltar.ripe
BLUP of genotypic value
Panel subpopulation or half-diallel cross
TE (n=61)
WE (n=86)
WW (n=87)
SxPN (n=53)
CSxPN (n=60)
GxPN (n=58)
TNxPN (n=47)
CSxS (n=60)
GxS (n=56)
TNxS (n=46)
GxCS (n=58)
TNxCS (n=55)
TNxG (n=52)
verday
BLUP of genotypic value
Panel subpopulation or half-diallel cross
TE (n=67)
WE (n=87)
WW (n=91)
SxPN (n=56)
CSxPN (n=65)
GxPN (n=63)
TNxPN (n=56)
CSxS (n=64)
GxS (n=59)
TNxS (n=62)
GxCS (n=63)
TNxCS (n=61)
TNxG (n=67)
samplday
BLUP of genotypic value
Panel subpopulation or half-diallel cross
TE (n=72)
WE (n=87)
WW (n=91)
SxPN (n=56)
CSxPN (n=65)
GxPN (n=63)
TNxPN (n=53)
CSxS (n=63)
GxS (n=59)
TNxS (n=59)
GxCS (n=63)
TNxCS (n=61)
TNxG (n=67)

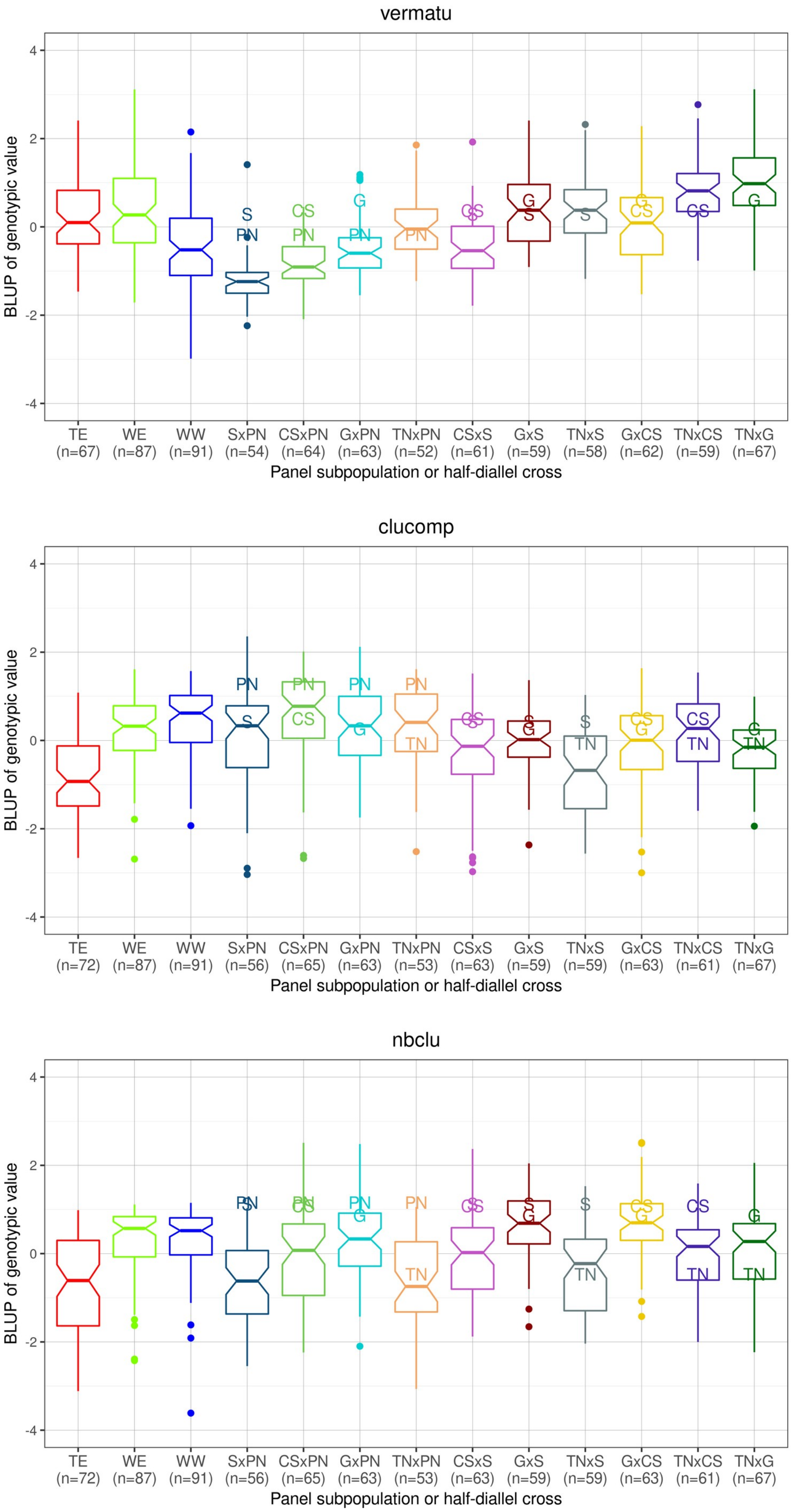
vermatu
BLUP of genotypic value
TE (n=67) WE (n=87) WW (n=91) SxPN (n=54) CSxPN (n=64) GxPN (n=63) TNxPN (n=52) CSxS (n=61) GxS (n=59) TNxS (n=58) GxCS (n=62) TNxCS (n=59) TNxG (n=67)
Panel subpopulation or half-diallel cross
clucomp
BLUP of genotypic value
TE (n=72) WE (n=87) WW (n=91) SxPN (n=56) CSxPN (n=65) GxPN (n=63) TNxPN (n=53) CSxS (n=63) GxS (n=59) TNxS (n=59) GxCS (n=63) TNxCS (n=61) TNxG (n=67)
Panel subpopulation or half-diallel cross
nbclu
BLUP of genotypic value
TE (n=72) WE (n=87) WW (n=91) SxPN (n=56) CSxPN (n=65) GxPN (n=63) TNxPN (n=53) CSxS (n=63) GxS (n=59) TNxS (n=59) GxCS (n=63) TNxCS (n=61) TNxG (n=67)
Panel subpopulation or half-diallel cross

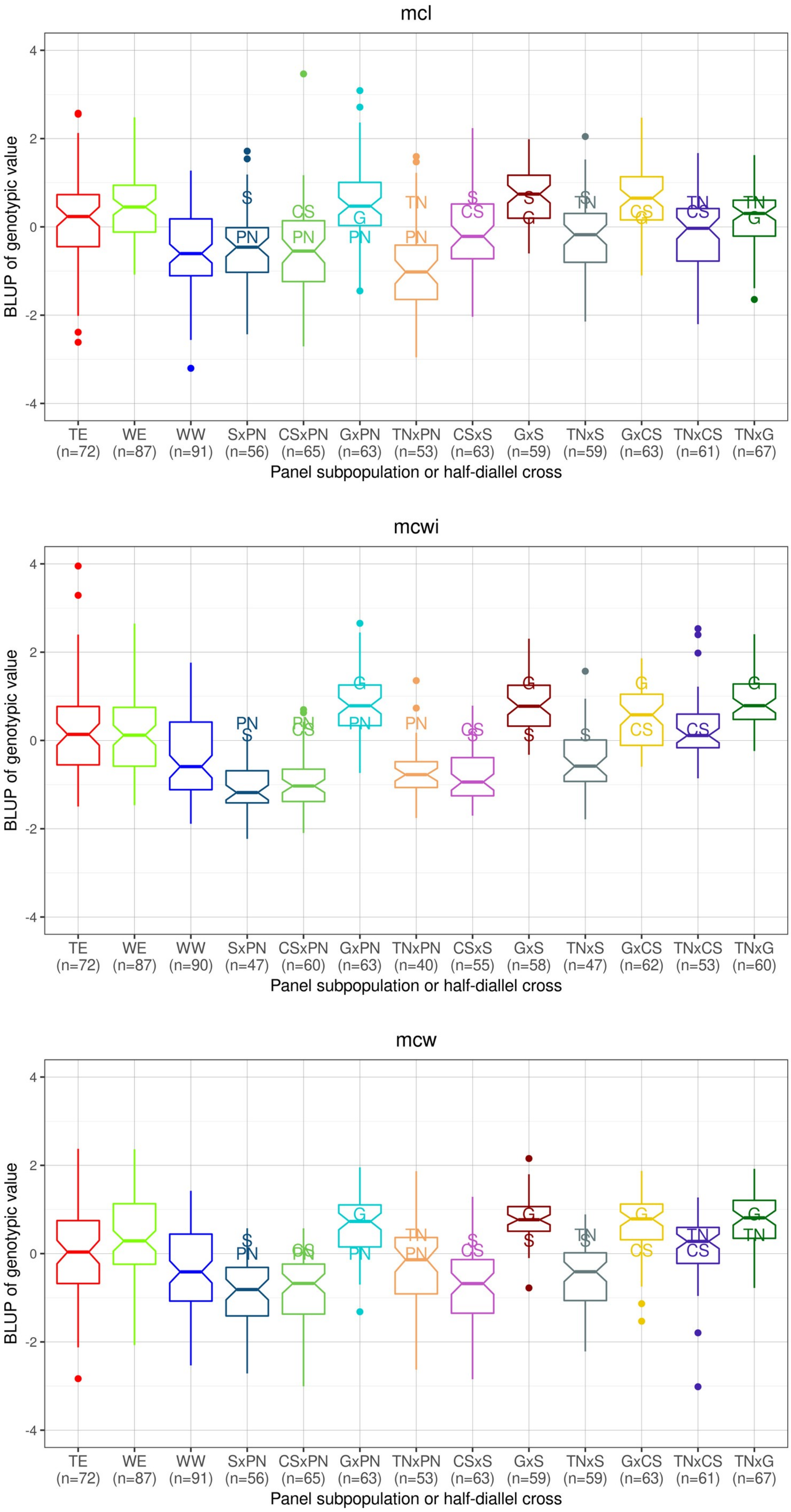
mcl
mcwi
mcw
BLUP of genotypic value
Panel subpopulation or half-diallel cross
TE (n=72) WE (n=87) WW (n=91) SxPN (n=56) CSxPN (n=65) GxPN (n=63) TNxPN (n=53) CSxS (n=63) GxS (n=59) TNxS (n=59) GxCS (n=63) TNxCS (n=61) TNxG (n=67)
TE (n=72) WE (n=87) WW (n=90) SxPN (n=47) CSxPN (n=60) GxPN (n=63) TNxPN (n=40) CSxS (n=55) GxS (n=58) TNxS (n=47) GxCS (n=62) TNxCS (n=53) TNxG (n=60)
TE (n=72) WE (n=87) WW (n=91) SxPN (n=56) CSxPN (n=65) GxPN (n=63) TNxPN (n=53) CSxS (n=63) GxS (n=59) TNxS (n=59) GxCS (n=63) TNxCS (n=61) TNxG (n=67)

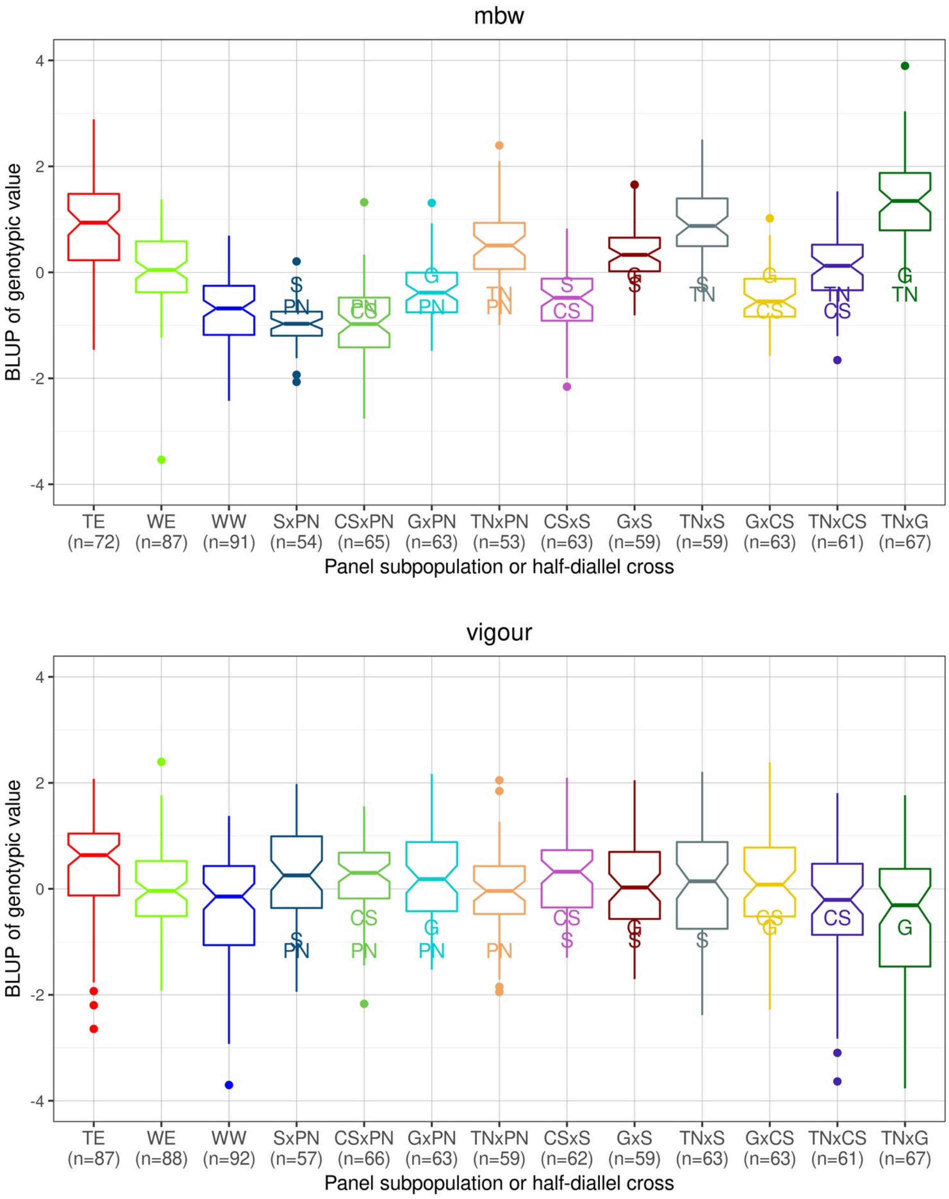


### Figure S3 Distribution of genotypic value estimates (BLUPs) for 15 traits, in each diversity panel subpopulation and each half-diallel cross.

Subpopulation or cross size is indicated below its name. All BLUPs were centered and scaled separately for each subpopulation or cross. TE: Table East, WE: Wine East, WW: Wine West. For traits, see abbreviations meaning in Methods section. Values of the parents are indicated by their initials as defined in **Table S4**.

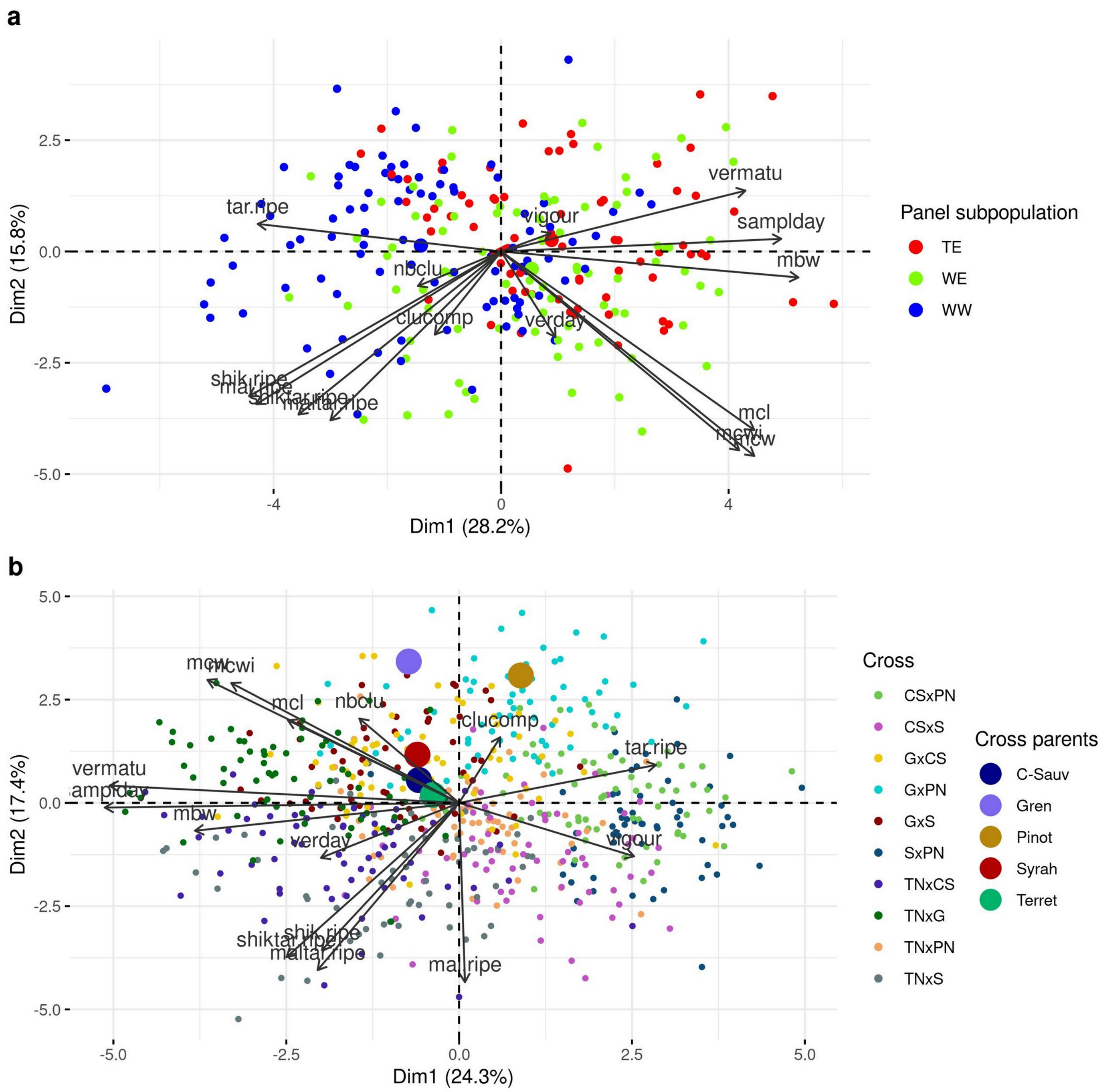


### Figure S4 PCA applied to genotypic BLUPs for the 15 traits.

a: In the diversity panel population (TE: Table East, WE: Wine East, WW: Wine West); b: In the half-diallel population. For traits, see abbreviations meaning in Methods section.

| **Parent 1 genotype** | **Parent 2 genotype** | **Progeny genotypes** (expected proportion in parenthesis) | **Parental average genotype** |
|---|---|---|---|
| **0** | **0** | **0** (1) | **0** |
| **0** | **1** | **0** (0.5) / **1** (0.5) | **0.5** |
| **0** | **2** | **1** (1) | **1** |

| | | | |
|---|---|---|---|
| **1** | **1** | **0** (0.25) / **1** (0.5) / **2** (0.25) | **1** |
| **1** | **2** | **1** (0.5) / **2** (0.5) | **1.5** |
| **2** | **2** | **2** (1) | **2** |

### Table S2 Computation of parental average genotypes.

For each cross and locus, parent genotypes (0, 1 or 2 for each parent) were used to derive the expected proportion of each genotype in the progeny under Mendelian segregation and from there, the parental average genotype. Probabilities are indicated in parentheses.

**a:**

| Cross | S1a-RR | S1a-LASSO | S1b-RR | S1b-LASSO | S2-RR | S2-LASSO |
|---|---|---|---|---|---|---|
| **CSxPN** | **0.33** | 0.03 | **0.4** | 0.38 | 0.22 | **0.35** |
| **CSxS** | **0.68** | 0.22 | 0.67 | **0.74** | **0.72** | 0.32 |
| **GxCS** | **0.52** | 0.43 | 0.42 | **0.45** | **-0.08** | -0.23 |
| **GxPN** | **0.6** | 0.52 | **0.56** | 0.49 | -0.45 | **-0.3** |
| **GxS** | **0.77** | 0.58 | **0.71** | 0.65 | 0 | **0.11** |
| **SxPN** | **0.29** | 0.06 | 0.37 | **0.42** | **0.61** | 0.35 |
| **TNxCS** | **0.57** | 0.15 | **0.61** | 0.59 | 0.41 | **0.63** |
| **TNxG** | **0.72** | 0.52 | **0.85** | 0.81 | 0.21 | **0.61** |
| **TNxPN** | **0.38** | 0.22 | **0.67** | 0.56 | -0.02 | **0.01** |
| **TNxS** | **0.78** | 0.36 | **0.78** | 0.77 | **0.25** | 0.22 |

**b:**

| Trait | S1a-RR | S1a-LASSO | S1b-RR | S1b-LASSO | S2-RR | S2-LASSO |
|---|---|---|---|---|---|---|
| **mal.ripe** | **0.9** | 0.82 | **0.91** | 0.82 | **0.73** | -0.01 |
| **tar.ripe** | **0.95** | 0.94 | **0.95** | 0.91 | **0.7** | 0.65 |
| **shik.ripe** | 1 | 1 | 0.98 | **0.99** | 0.19 | **0.96** |
| **shiktar.ripe** | **1** | 0.99 | 0.98 | **1** | 0.16 | **0.56** |
| **maltar.ripe** | **0.98** | 0.96 | 0.96 | **0.97** | **0.72** | 0.03 |
| **verday** | 0.93 | **0.95** | **0.92** | 0.9 | 0.66 | **0.81** |
| **sampldday** | 0.99 | 0.99 | 0.98 | 0.98 | **0.82** | 0.59 |
| **vermatu** | **0.97** | 0.95 | 0.95 | 0.95 | 0.63 | **0.72** |
| **clucomp** | **0.84** | 0.55 | **0.87** | 0.75 | 0.26 | **0.55** |
| **nbclu** | 0.82 | **0.91** | **0.95** | 0.91 | **-0.64** | -0.75 |
| **mcl** | 0.85 | **0.93** | 0.86 | **0.92** | 0.52 | **0.76** |
| **mcwi** | 0.94 | 0.94 | **0.94** | 0.92 | 0.77 | **0.83** |
| **mcw** | **0.93** | 0.79 | 0.96 | 0.96 | 0.74 | **0.81** |
| **mbw** | **0.96** | 0.94 | 0.96 | **0.97** | **0.83** | 0.63 |
| **vigour** | **0.85** | 0.67 | **0.86** | 0.77 | -0.15 | **-0.09** |

### Table S3 Predictive ability of cross mean.

Pearson's correlation between the observed cross mean and the one predicted based on parental average genotypes. Values are reported for scenarios 1a, 1b and 2, with RR and LASSO methods. The best PA value for each scenario is indicated in bold.

**a**: per-cross PA (correlation based on 15 observations); **b** per-trait PA (correlation based on 10 observations).

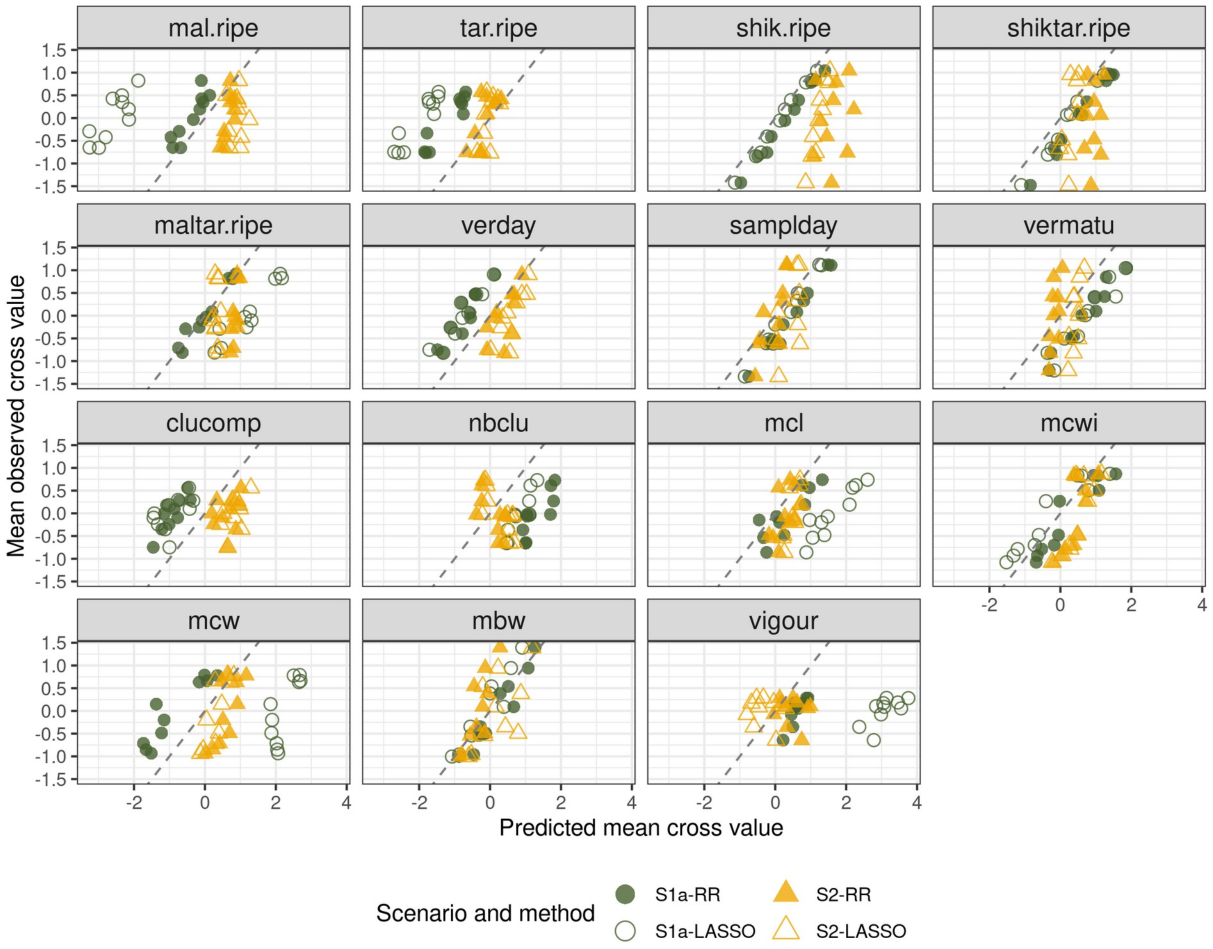


### Figure S5 Observed vs predicted cross means for each trait in the half-diallel

Observed vs predicted (based on parental average genotypes) cross mean for each trait in the half-diallel, according to four prediction modalities: with allelic effects estimated in the half-diallel or in the diversity panel (*S1a and S2*, respectively) and with RR or the LASSO. The dashed line indicates the perfect fit (with slope=1 and intercept=0), points deviating from this line indicate bias.

mal.ripe

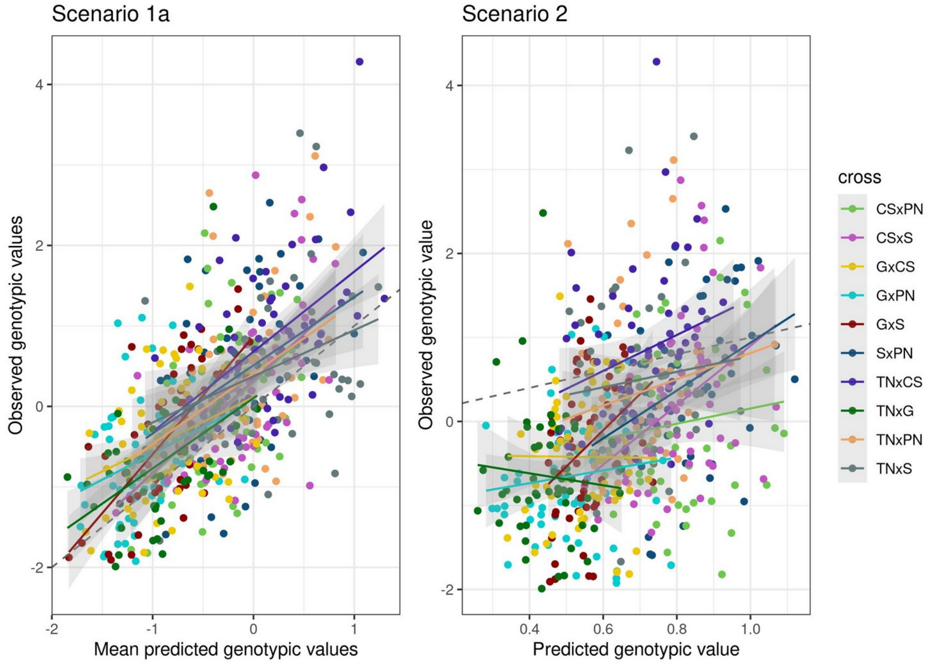


tar.ripe

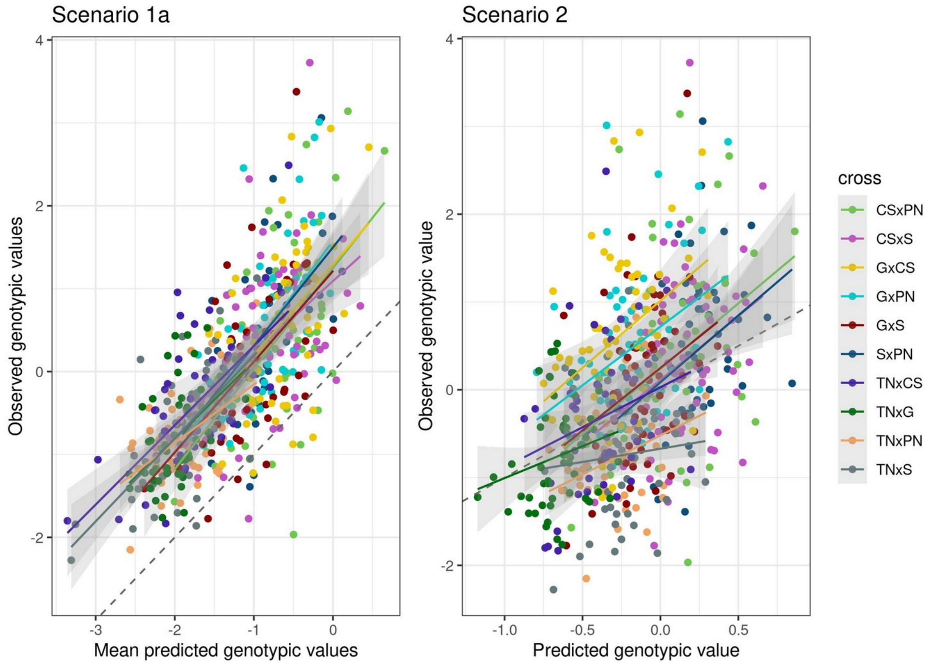

shik.ripe

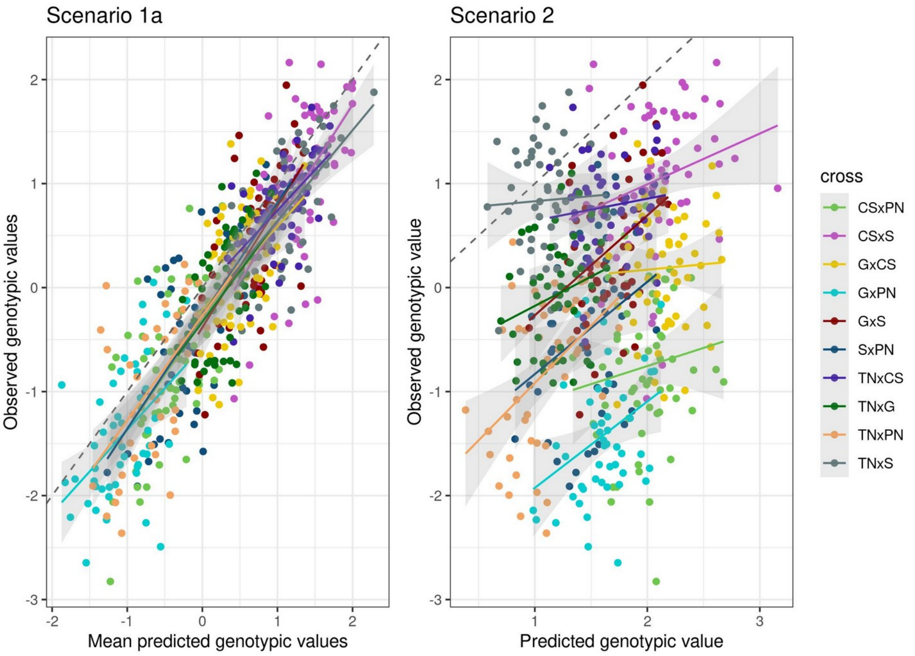


shiktar.ripe

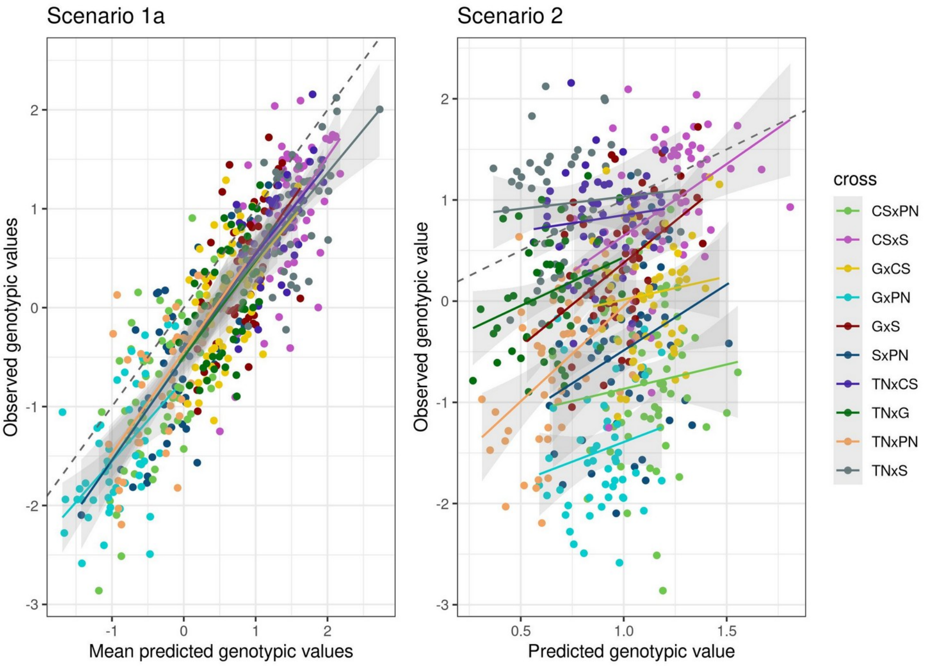

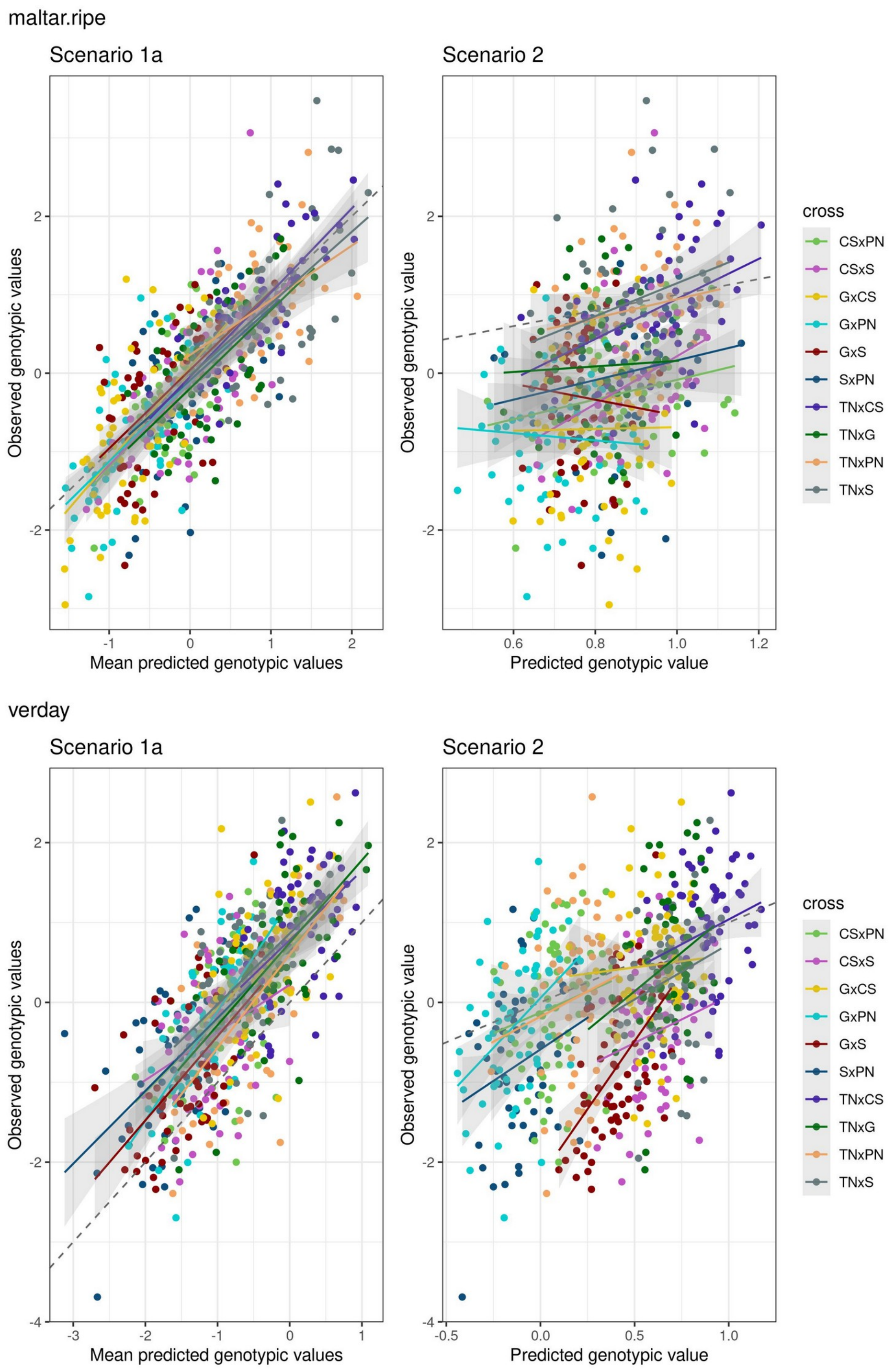
maltar.ripe
Scenario 1a
Scenario 2
Observed genotypic values
Mean predicted genotypic values
Observed genotypic value
Predicted genotypic value
cross
CSxPN
CSxS
GxCS
GxPN
GxS
SxPN
TNxCS
TNxG
TNxPN
TNxS
verday
Scenario 1a
Scenario 2
Observed genotypic values
Mean predicted genotypic values
Observed genotypic value
Predicted genotypic value
cross
CSxPN
CSxS
GxCS
GxPN
GxS
SxPN
TNxCS
TNxG
TNxPN
TNxS

sampIday

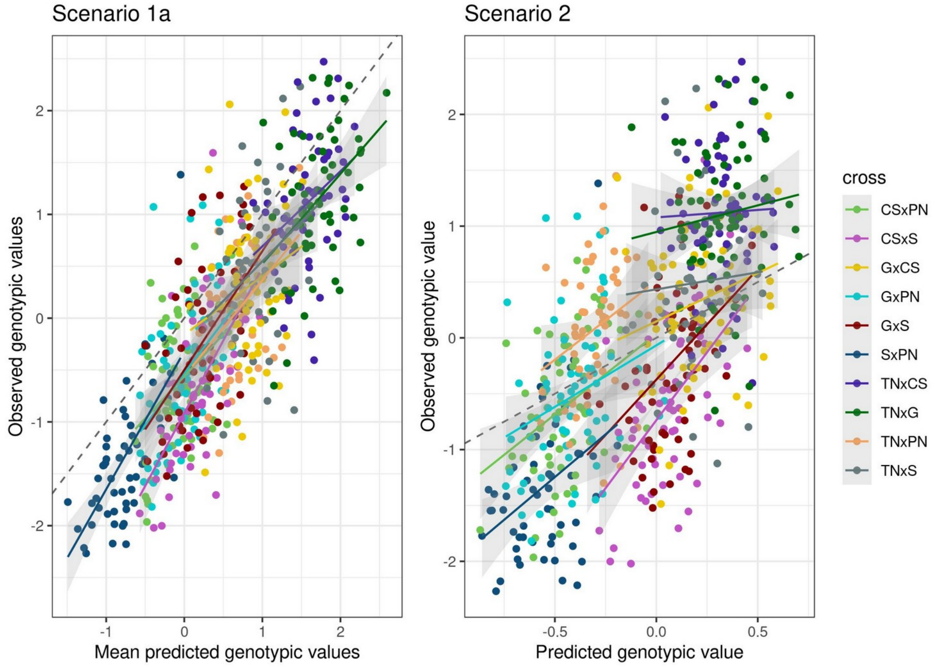
Scenario 1a
Scenario 2
Observed genotypic values
Mean predicted genotypic values
Observed genotypic value
Predicted genotypic value
cross
CSxPN
CSxS
GxCS
GxPN
GxS
SxPN
TNxCS
TNxG
TNxPN
TNxS

vermatu

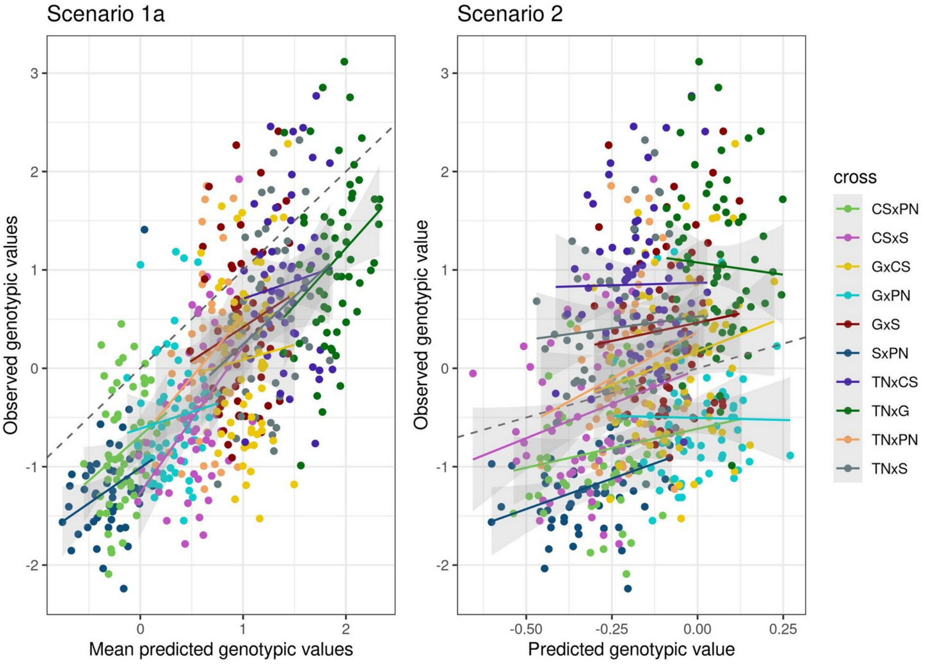


clucomp

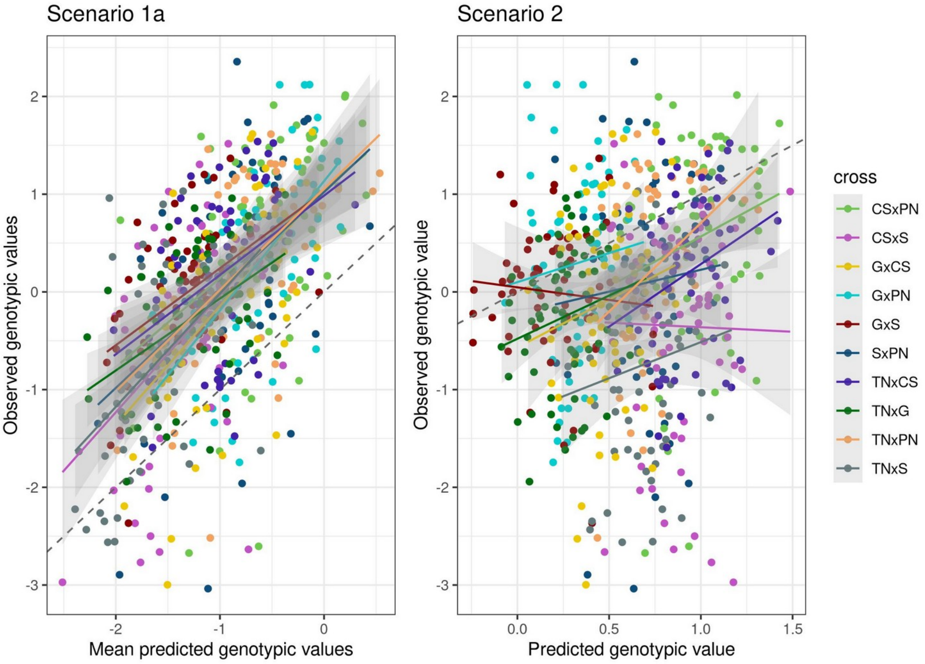

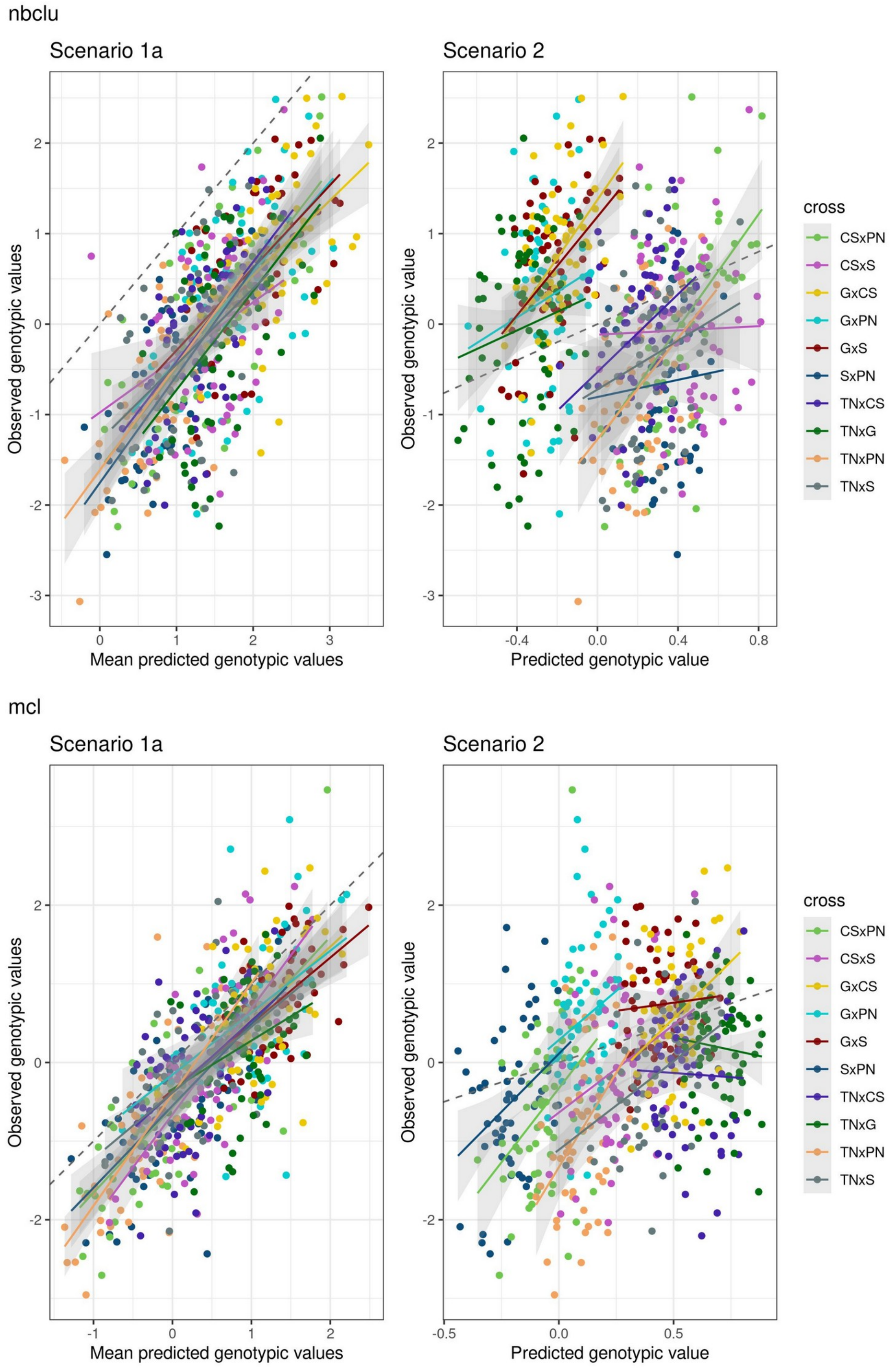
nbclu
Scenario 1a
Scenario 2
Observed genotypic values
Mean predicted genotypic values
Observed genotypic value
Predicted genotypic value
cross
CSxPN
CSxS
GxCS
GxPN
GxS
SxPN
TNxCS
TNxG
TNxPN
TNxS
mcl
Scenario 1a
Scenario 2
Observed genotypic values
Mean predicted genotypic values
Observed genotypic value
Predicted genotypic value
cross
CSxPN
CSxS
GxCS
GxPN
GxS
SxPN
TNxCS
TNxG
TNxPN
TNxS

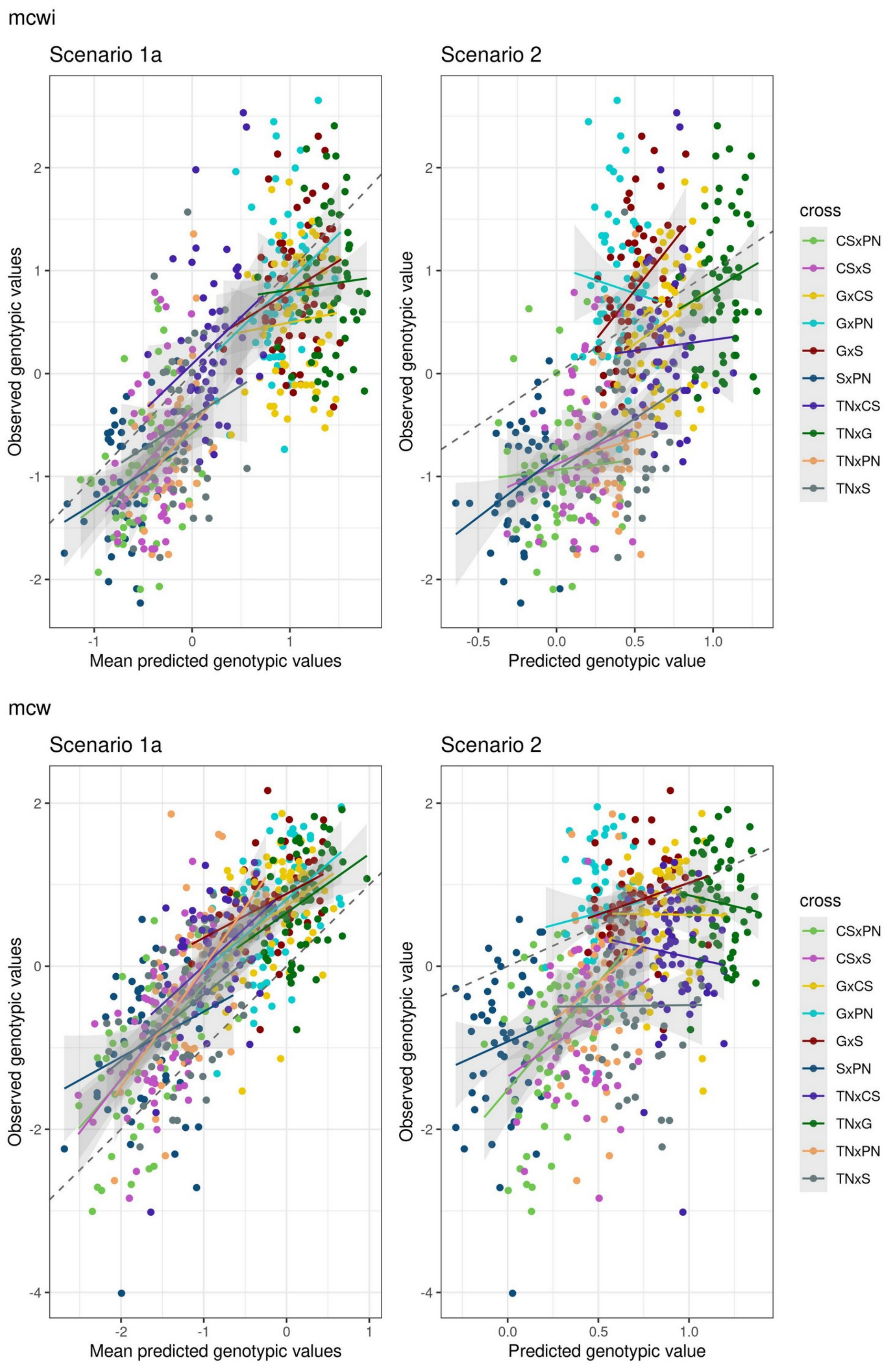
mcwi
Scenario 1a
Scenario 2
Observed genotypic values
Mean predicted genotypic values
Observed genotypic value
Predicted genotypic value
cross
CSxPN
CSxS
GxCS
GxPN
GxS
SxPN
TNxCS
TNxG
TNxPN
TNxS
mcw
Scenario 1a
Scenario 2
Observed genotypic values
Mean predicted genotypic values
Observed genotypic value
Predicted genotypic value
cross
CSxPN
CSxS
GxCS
GxPN
GxS
SxPN
TNxCS
TNxG
TNxPN
TNxS

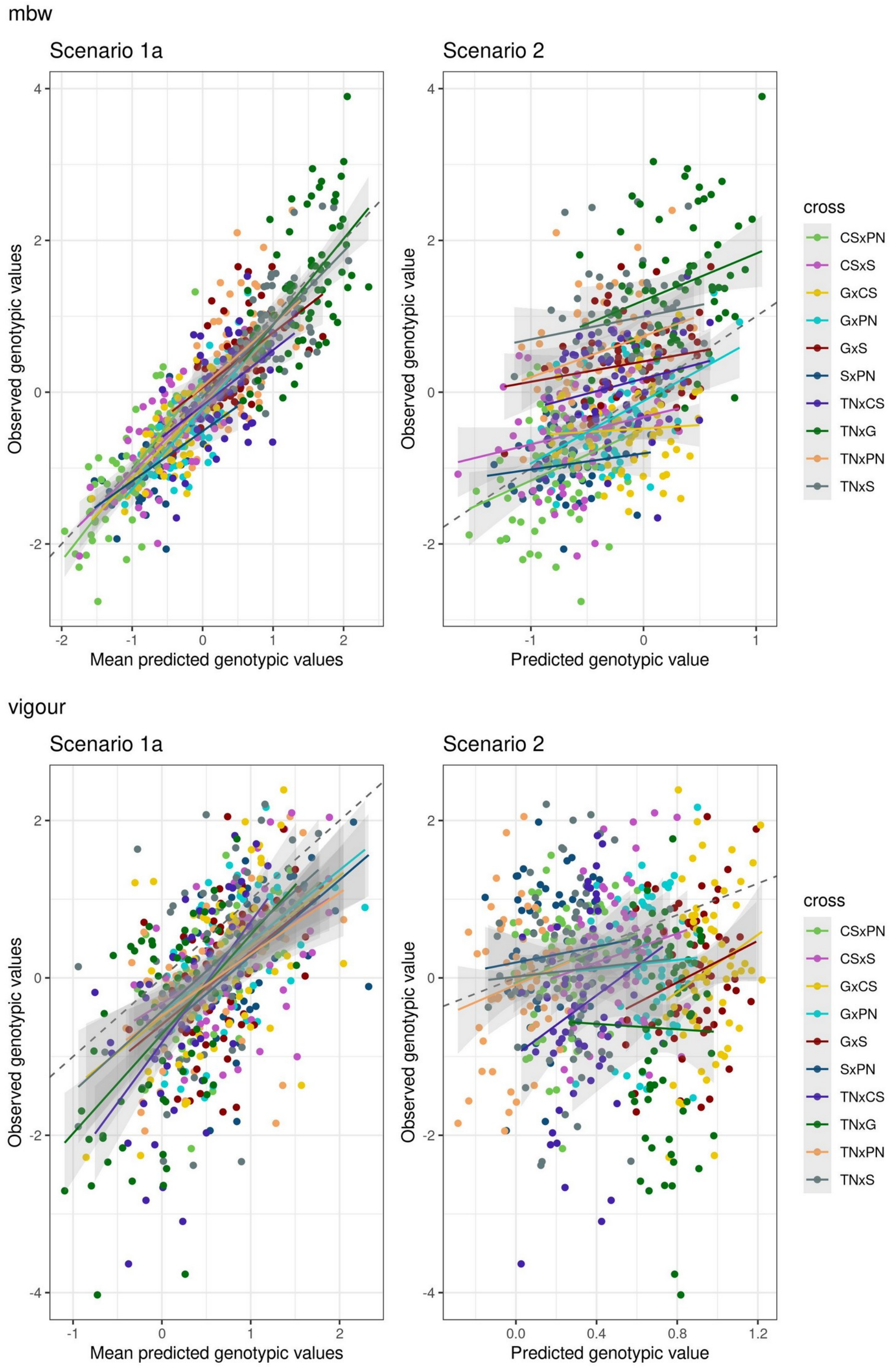


### Figure S6 Observed vs predicted individual genotypic values for 15 traits.

Comparison between scenarios 1a (left) and 2 (right). Each point represents one offspring of one cross. A linear regression was fitted for each cross, with standard error displayed in grey. For scenario 1a, predicted genotypic values were averaged over 10 cross-validation repetitions. Identity (y=x) is displayed with a dashed line. Genotypic values were predicted using RR method.

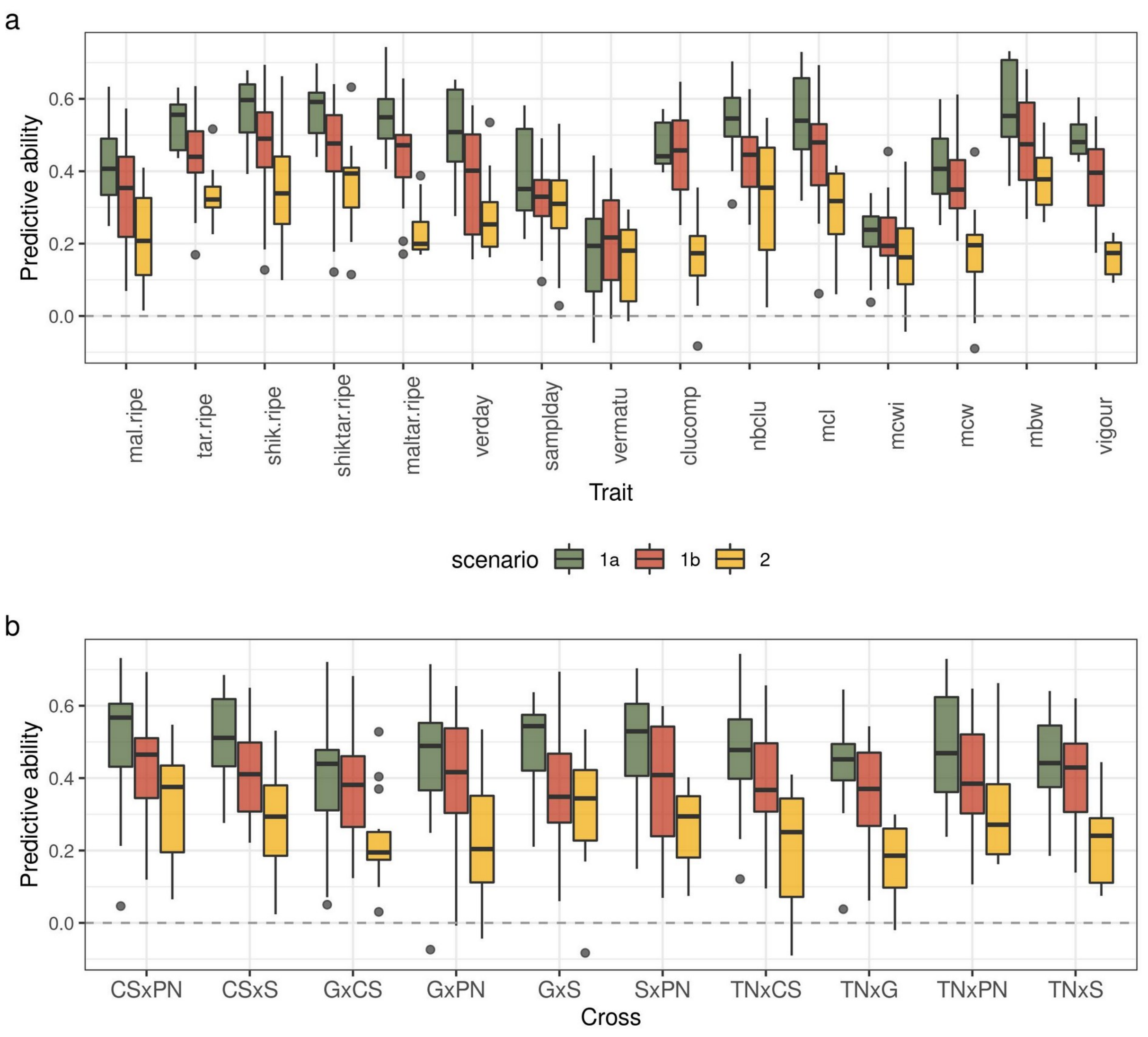


### Figure S7 Distribution of predictive ability for Mendelian sampling genomic prediction.

Boxplots of PA values, calculated for all individuals within a cross, for scenarios 1a (green), 1b (red) and 2 (yellow), for the best method among RR and LASSO

**a**: distribution for each trait, **b**: distribution for each cross,

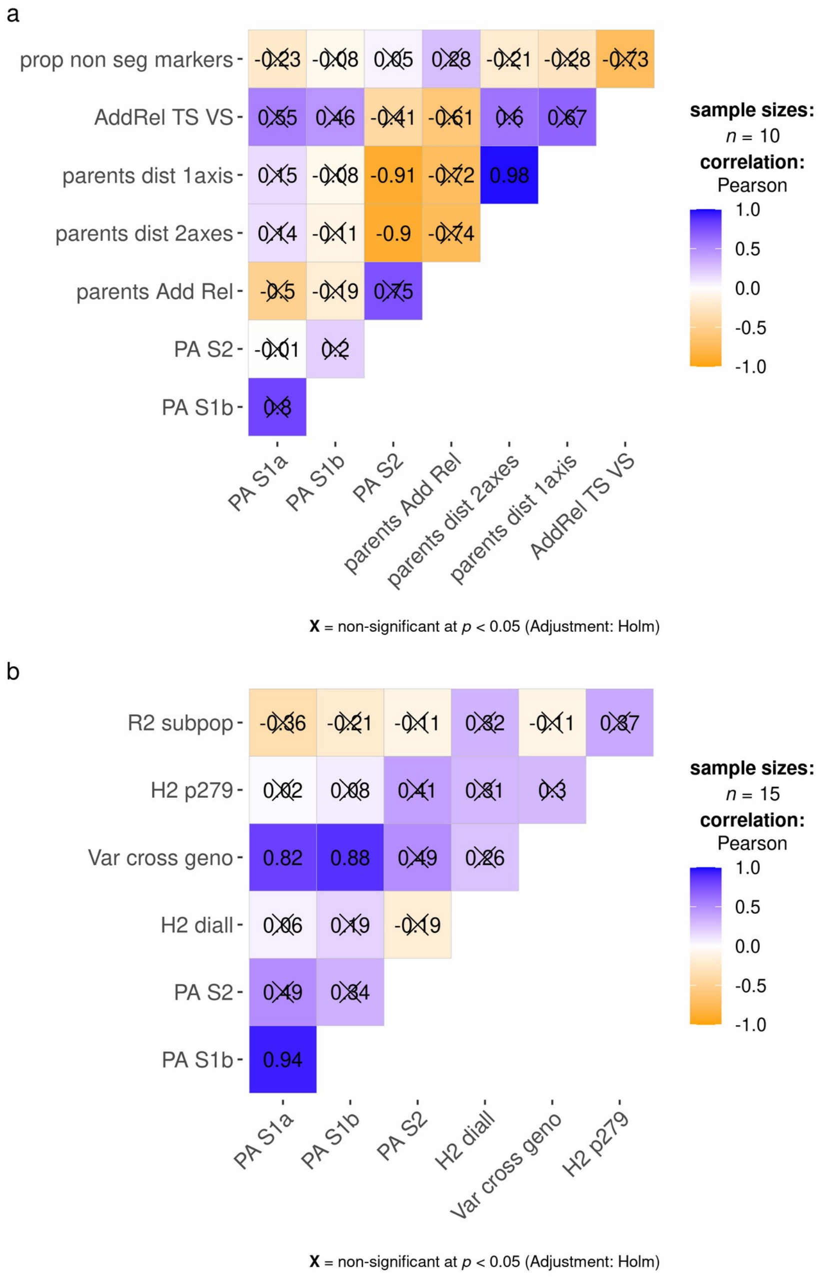


### Figure S8 Correlation plot for PA of cross mean and potential explanatory variables.

PA S1a, PA S1b, PA S2 are PAs for the three scenarios 1a, 1b and 2, respectively.
**a**: per-cross PA and genetic variables. Prop non seg markers is the proportion of non-segregating markers in each half-diallel cross. AddRel TS VS is the mean additive relationship between TS and VS. Parents dist 1axis and parents dist 2axes are the pairwise distances between half-diallel parents on the PCA for the first axis or the first two axes, respectively. Parents Add Rel is the pairwise additive relationship between half-diallel parents.

**b**: per-trait PA and trait-related variables. H2 diall and H2 p279 are broad-sense heritability in the half-diallel and the diversity panel, respectively. Var cross geno is the proportion of genetic variance due to differences between crosses, as described in Methods.

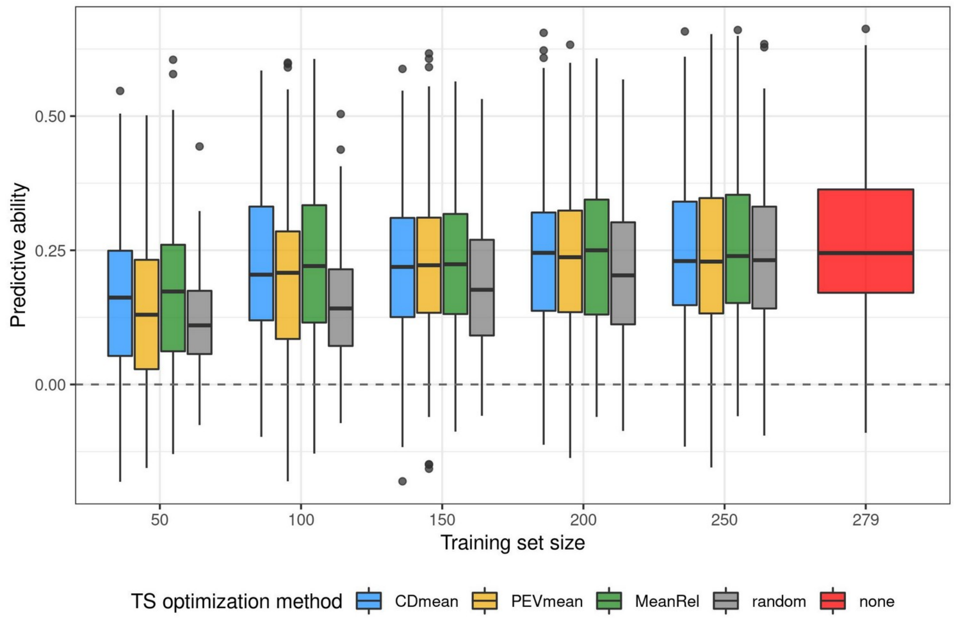


### Figure S9 Distribution of predictive ability for Mendelian sampling genomic prediction, after training set optimization.

Boxplots of individual PA values over all traits and crosses in scenario 2, for different TS sizes and optimization methods, as described in Methods. "Random" method corresponds to random sampling of TS genotypes. "None" corresponds to the use of the whole diversity panel as TS. Optimization was performed for each cross of the half-diallel. The best predictive ability was kept between RR and LASSO for each trait and cross.

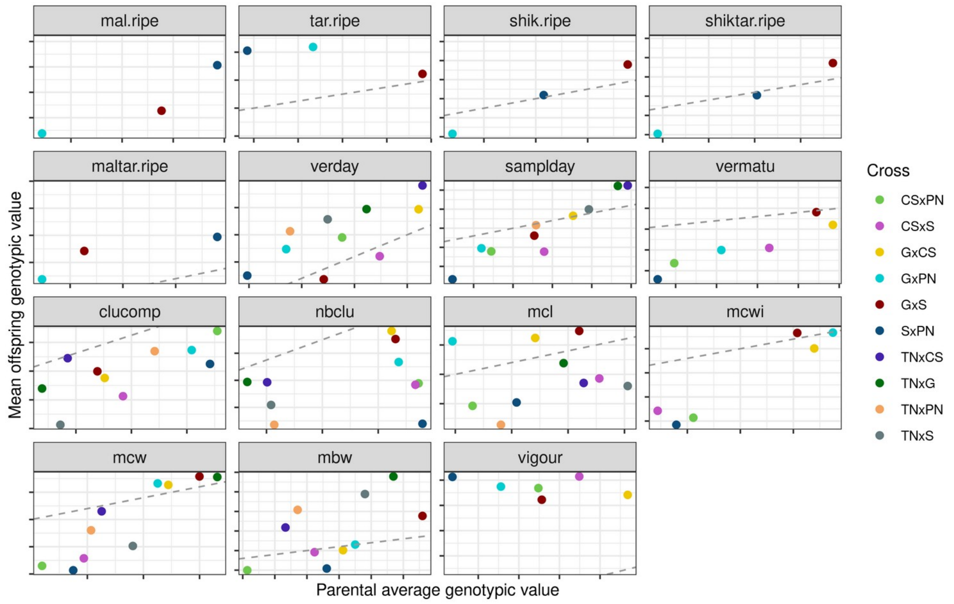


### Figure S10 Mean offspring observed genotypic value vs parental average observed genotypic value in each half-diallel cross for 15 traits

The grey dashed line corresponds to identity (x=y).

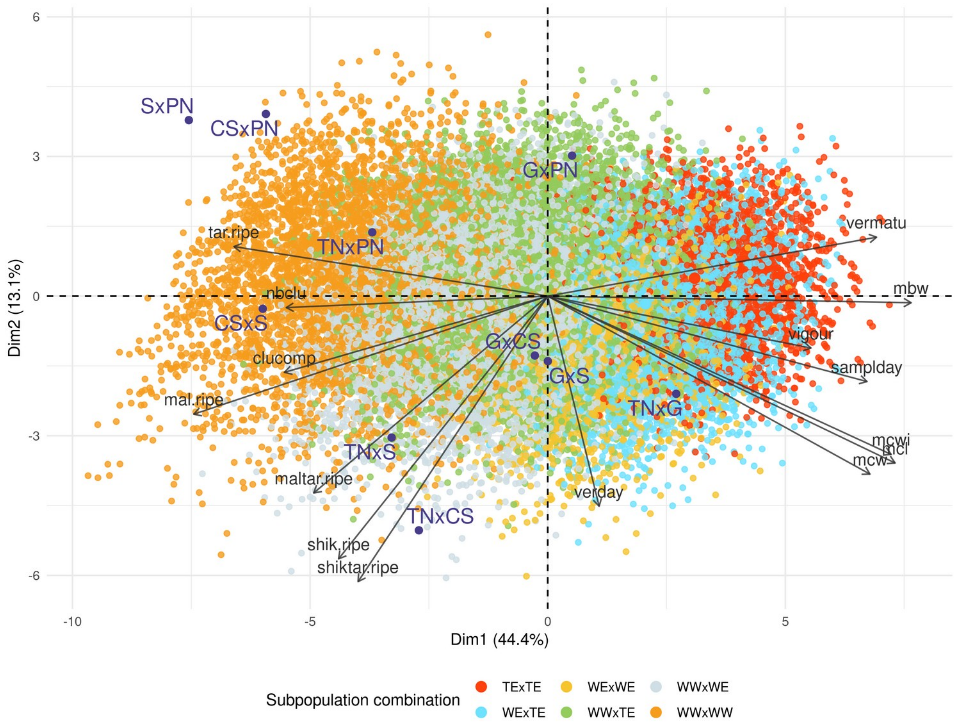


### Figure S11 PCA of predicted cross mean genotypic values for all 38,781 possible simulated crosses between the 279 varieties of the diversity panel

Prediction was based on parental average genotypes and marker effects estimated with RR in the diversity panel. For each simulated cross, the dot color corresponds to the combination of panel subpopulations from which the parents of the cross originate. Values of the half-diallel crosses were projected.

| Cross | Female | Male |
|---|---|---|
| SxPN | NA | PN: Pinot Noir |
| CSxPN | CS: Cabernet-Sauvignon | PN: Pinot Noir |
| GxPN | G: Grenache | PN: Pinot Noir |
| TNxPN | NA | PN: Pinot Noir |
| CSxS | CS: Cabernet-Sauvignon | S: Syrah |
| GxS | G: Grenache | S: Syrah |
| TNxS | TN: Terret Noir | S: Syrah |
| GxCS | G: Grenache | NA |
| TNxCS | TN: Terret Noir | NA |
| TNxG | TN: Terret Noir | NA |

### Table S4 Partial pedigree of half-diallel crosses used for marker imputation.

As the software **Fimpute3** does not handle hermaphroditism, we declared a partial pedigree which maximizes the number of crosses with both parents defined.

| | Training set | Training set sizes (depending on trait) | Validation set | Validation set sizes (depending on trait) |
|---|---|---|---|---|
| Scenario 1a | Random sampling of 9/10th of the half-diallel | min: 477<br>max: 558 | The remaining 1/10th of the half-diallel | min: 53<br>max: 62 |
| Scenario 1b | Three bi-parental crosses with one common parent | min: 140<br>max: 193 | The fourth cross with the same common parent | min: 40<br>max: 67 |
| Scenario 2 | The whole diversity panel | min: 234<br>max: 267 | Each cross of the half-diallel | min: 531<br>max: 620 |

#### Table S5 Training and validation sets composition for each scenario used to assess genomic prediction

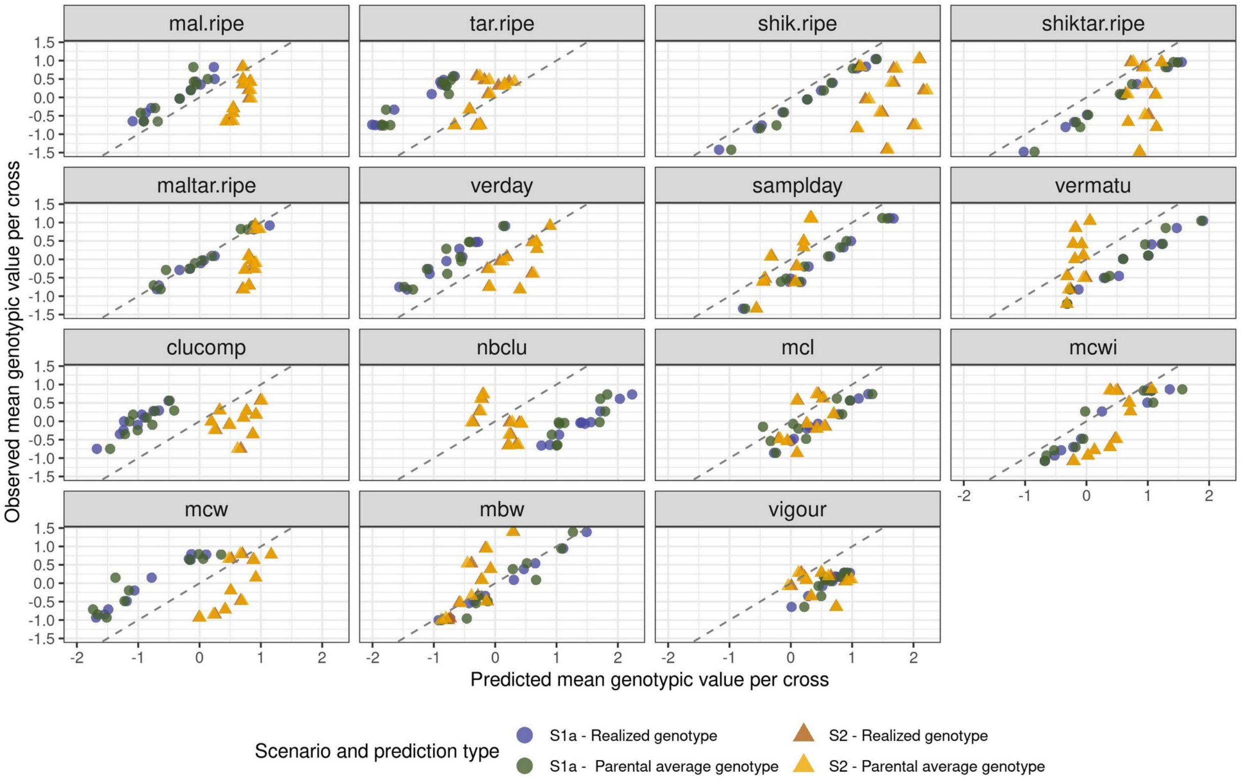


#### Figure S12 Observed vs predicted cross mean for 15 traits.

Predicted values per cross are displayed for realized genotypes or parental average genotype, and for scenarios 1a (circle) or 2 (triangle). Observed cross mean is the averaged genotypic value over all offspring within a cross. Identity (y=x) is displayed with a dashed line. Predicted values were obtained with RR.
